# Engineered probiotics for tumor-targeted combination chemoimmunotherapy

**DOI:** 10.64898/2026.02.04.703875

**Authors:** Zaofeng Yang, Jongwon Im, Noah Chen, Dylan L. Mariuzza, Kenia de los Santos-Alexis, Fangda Li, Tal Danino, Nicholas Arpaia

**Affiliations:** Department of Microbiology and Immunology, Columbia University, New York, NY 10032, USA; Department of Biomedical Engineering, Columbia University, New York, NY 10027, USA; Herbert Irving Comprehensive Cancer Center, Columbia University, New York, NY 10032, USA; Data Science Institute, Columbia University, New York, NY 10027, USA

## Abstract

Achieving tumor-specific delivery and sustained activation of both cytotoxic and immune-modulating agents remains a critical challenge in chemoimmunotherapy. Here, we present a bacterial platform engineered to combine enzyme/prodrug chemotherapy with immunotherapy, where tumor-homing *E. coli* Nissle 1917 expresses cytosine deaminase to convert the prodrug 5-fluorocytosine into the cytotoxic drug 5-fluorouracil within tumors. Concurrently, the engineered bacteria produce an IL-15 superagonist and a PD-L1 blocking nanobody to mitigate the immunosuppressive effects of tumor-localized chemotherapy. This platform demonstrated potent antitumor effects in the murine MC38 solid tumor model. Mechanistic studies showed that the combination therapy enhances activation of antigen-presenting cells, T cells and natural killer cells, while reducing immunosuppressive populations. In summary, our approach integrates enzyme/prodrug therapy and immunotherapy into a single bacterial delivery system, overcoming the limitations of conventional therapies and offering a scalable and precision-engineered strategy with an improved safety profile for synergistic cancer treatment.

**One Sentence Summary:** A precision-engineered bacterial platform integrates enzyme/prodrug chemotherapy and immunotherapy to drive synergistic antitumor responses with enhanced safety, offering promise for clinical translation.

## INTRODUCTION

Chemotherapy, a cornerstone of cancer treatment, employs cytotoxic drugs to kill rapidly dividing tumor cells (*1*). However, its lack of specificity often results in collateral damage to healthy cells, leading to significant side effects (*1*). One promising strategy for targeting chemotherapeutics to tumors is enzyme/prodrug therapy (EPT) (*2, 3*). This two-step approach involves first delivering a catalytic enzyme to tumor tissues, followed by the systemic administration of a nontoxic prodrug. The prodrug is then converted into a cytotoxic drug *in situ* by the tumor-localized enzyme (*2, 3*). Cytosine deaminase (CD) and 5-fluorocytosine (5-FC) represent a classic example of EPT, where CD catalyzes the conversion of 5-FC to a commonly used anticancer drug 5-fluorouracil (5-FU) (*2–4*). Ideally, EPT would enable tumor-specific activation of anticancer prodrugs and is therefore expected to significantly reduce the off-target toxicities associated with conventional chemotherapeutics. Nevertheless, the key challenges for efficacious EPT lie in: (a) achieving tumor-selective delivery of the prodrug-converting enzyme, and (b) ensuring sustained expression or improved retention of the converting enzyme within tumor sites.

Another limitation of chemotherapy is that most chemotherapeutics exhibit low response rates when used as single agents (*1, 4*), largely due to tumor heterogeneity and the emergence of drug resistance, necessitating the development of new modalities and combination strategies (*4–6*). Immunotherapy, which harnesses the host’s immune system to combat tumors, has revolutionized cancer treatment in recent years (*7*). Immune checkpoint inhibitors (ICIs), such as antibodies targeting programmed cell death protein 1 (PD-1), and its ligand (PD-L1), can relieve T cells from immunosuppression, enabling them to eliminate tumor cells (*8*). Despite promising antitumor responses across a broad spectrum of cancers, the efficacy of ICIs is heavily dependent on tumor immunogenicity, with response rates ranging from 15% to 30% in most solid tumors (*8, 9*). Of note, numerous studies demonstrate that conventional chemotherapies exert complex immunomodulatory effects (*10*). Many chemotherapeutics can promote antitumor immune responses by increasing the immunogenicity of malignant cells or by inhibiting immunosuppressive cell populations, while some may also impair such responses by depleting effector cells or indirectly provoking anergy (*10–12*). For example, 5-FU facilitates tumor antigen uptake by dendritic cells and induces T cell immunity for tumor control (*11, 13*), but it simultaneously upregulates co-inhibitory molecules such as PD-L1 on tumor cells or tumor-infiltrating myeloid cells, which suppress T cell functionality (*12*). Given these facts, considerable efforts have been directed toward combining immunotherapy with chemotherapy to improve disease control in two scenarios: (a) maximizing the immunostimulatory effects of chemotherapies, and (b) mitigating their immunosuppressive effects (*10, 12*). Although both preclinical and clinical results have highlighted the notable potential of chemoimmunotherapy combinations (*14–16*), designing rational combinations with optimal efficacy and safety remains challenging (*17*); specifically: (a) a deeper understanding of the impact chemotherapies have on the immune response is required to prioritize the myriad possible combinations, and (b) because immunotherapy can often lead to moderate-to-severe immune-related adverse events (*12*), combining it with chemotherapy may increase overall toxicity (*16*).

To overcome these obstacles, we have developed a novel combination strategy that integrates EPT with immunotherapy through a single bacterial platform. One appealing feature of bacteria is their ability to selectively colonize tumor cores and preferentially expand within the immune-privileged tumor microenvironment (TME) (*18, 19*). Furthermore, synthetic biology approaches enable the programming of bacteria to produce therapeutic proteins, such as chemokines and nanobodies, for tumor-targeted delivery (*20–23*). By leveraging these advantages, we program bacteria to express CD for sustained enzyme production within tumor tissues, resulting in efficient CD/5-FC EPT with enhanced safety. More importantly, mechanistic studies on how bacteria-based EPT reshapes tumor-infiltrating immune cells direct us to further engineer bacteria to co-express an IL-15 superagonist and a PD-L1 blocking nanobody alongside CD, culminating in a rational chemoimmunotherapy combination elicited by a single engineered bacterial vector (**Fig. 1A**). We demonstrate that a single dose of these engineered bacteria, followed by prodrug administration, effectively activates intratumoral antigen-presenting cells (APCs), CD4 T helper cells, and tumoricidal CD8 T cells and NK cells. This widespread activation of immune cells leads to a remarkable enhancement in therapeutic efficacy compared to EPT alone. Unlike traditional combination therapies, which often face challenges in achieving precise tumor targeting and durable therapeutic efficacy, this microbial chemoimmunotherapy induces local antitumor immune responses, all while minimizing toxicity.

**Fig. 1.**
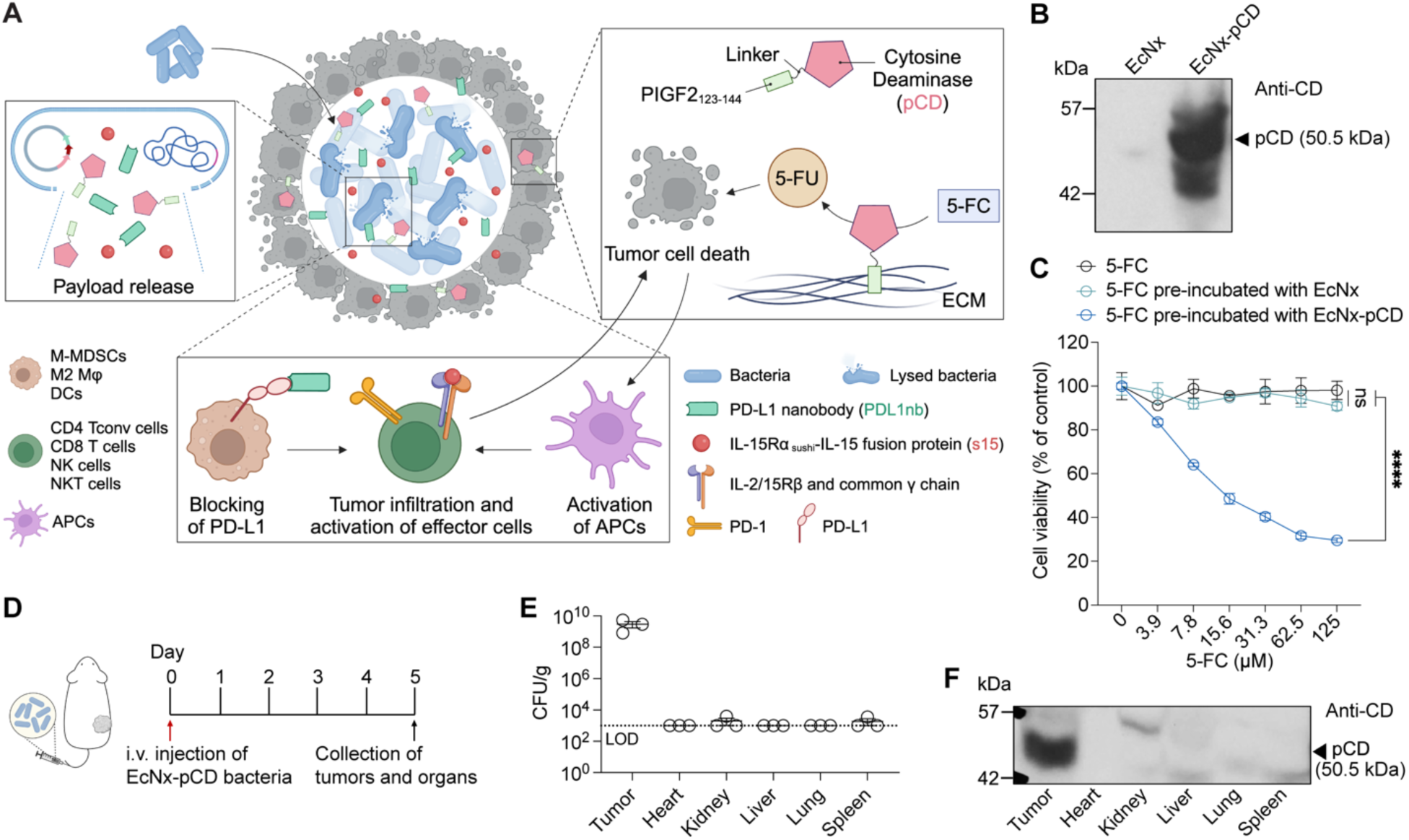
EcNx-pCD bacteria produce pCD exclusively in tumor tissues. **(A)** Schematic illustrating the bacterial platform for enzyme/prodrug therapy (EPT). Tumor-targeting *E. coli* Nissle 1917 produces and releases cytosine deaminase *in situ,* catalyzing the conversion of the nontoxic prodrug 5-FC into the cytotoxic agent 5-FU. Cytosine deaminase is tagged with a PIGF2_123-144_ peptide (pCD) that broadly binds to matrix proteins in the tumor microenvironment, restricting its diffusion beyond ECM-rich tumor margins. Additionally, the co-production of immune-modulating IL-15Rα_sushi_-IL15 fusion protein (s15) and PD-L1 nanobody (PDL1nb) augments antitumor immunity and synergizes with EPT to improve tumor control. ECM, extracellular matrix; M-MDSCs, monocytic myeloid-derived suppressor cells; Mφ, macrophages; DCs, dendritic cells; NK/NKT cells, natural killer/natural killer T cells; APCs, antigen-presenting cells. **(B)** Western blot image showing the expression of pCD by EcNx-pCD bacteria. **(C)** MC38 cell viability showing the activity of pCD. 2 mM 5-FC was pre-incubated with EcNx or EcNx-pCD bacteria in 1 mL DMEM (OD_600_ = 0.8) for 30 min. The bacteria were then pelleted down, and the supernatants were diluted and added to MC38 cells in a 96-well plate. Cell viability was determined using MTT after 48h incubation. Data are presented as mean ± sem. ns, not significant; **** p<0.0001; Two-way ANOVA with Holm-Šídák multiple comparisons test. **(D)** Schematic illustration of experimental procedures for evaluating the tumor targeting efficiency of EcNx-pCD bacteria. Mice bearing MC38 tumors were intravenously injected 2 × 10^7^ CFU/mouse EcNx-pCD bacteria. Tumors and organs were collected for CFU and pCD expression analyses 5 days post bacteria injection. **(E)** Colony-forming unit (CFU) analysis of different organs and tumors. Data are presented as mean ± sem. **(F)** Western blot image showing the tumor-exclusive expression of pCD protein.

## RESULTS

### EcNx-pCD bacteria target tumors and locally produce cytosine deaminase

We initiated our studies by evaluating the efficacy and toxicity of 5-FU using the syngeneic MC38 colorectal tumor model. While three intraperitoneal (i.p.) injections of 5-FU on day 10, 12 and 14 post-tumor cell inoculation slowed the growth of subcutaneous MC38 tumors, mice receiving 5-FU treatment showed significant body weight decrease (7% decrease on day 16), and had not fully recovered in the following 14 days when compared to the PBS control mice (**fig. S1**). Given the extensive use of 5-FU as a chemotherapy drug, along with the effectiveness and systemic toxicity demonstrated above and suggested by previous studies (*1, 24, 25*), we reasoned that the CD/5-FC pair could be a viable candidate for further exploration among enzyme/prodrug therapies.

To achieve tumor-specific conversion of prodrug 5-FC to cytotoxic 5-FU, EcNx—an engineered *Escherichia coli* Nissle 1917 (EcN) strain with an integrated lysing circuit for quorum-regulated bacterial cell lysis and release of genetically encoded payloads (*20, 26*)—was utilized as a vehicle for delivering prodrug-converting enzyme CD to tumors. We hypothesized that, following tumor-selective colonization, these bacteria would continuously produce CD within tumor tissues and subsequently lyse to release CD into the TME after reaching a quorum threshold. To further avoid potential diffusion of the released enzyme beyond ECM-dense tumor margins, a heparin-binding peptide derived from placenta growth factor 2 (PIGF2_123-144_)—which broadly anchors to the ECM contents of TME such as collagens, fibronectins, and heparan sulfate proteoglycans (HSPGs) (*19, 27*)—was genetically fused to the N-terminus of CD (herein termed, “pCD”) (*28*). Production of pCD by engineered bacteria was verified by western blot analysis of lysates from EcNx and pCD-expressing EcNx (EcNx-pCD) bacteria using an anti-CD antibody (**Fig. 1B**). The enzymatic activity of pCD was then evaluated by incubating 5-FC with EcNx or EcNx-pCD bacteria. The resultant supernatants after incubation and centrifugation were added to MC38 cells for cell viability assay. As shown in **Fig. 1C**, 5-FC pre-incubated with EcNx-pCD bacteria significantly suppressed MC38 cell growth in a dose-dependent manner, whereas fresh 5-FC or 5-FC pre-incubated with EcNx control bacteria had no effect on cell viability, indicating that the recombinant pCD successfully converted 5-FC to cytotoxic 5-FU.

We next sought to examine the tumor colonization and *in situ* production dynamics of pCD by EcNx-pCD bacteria. Mice bearing subcutaneous MC38 tumors received a single intravenous (i.v.) injection of EcNx-pCD bacteria, and 5 days later, tumor, heart, kidney, liver, lung, and spleen were collected to assess biodistribution and detect the presence of pCD in tissues (**Fig. 1D**). The bacterial load in all major organs was below or near the limit of detection (**Fig. 1E**). In contrast, the average bacterial count in tumor tissues exceeded 10^9^ CFU/g (**Fig. 1E**), suggesting that EcNx-pCD bacteria persisted within tumors while being rapidly cleared from other organs. Consistent with biodistribution analysis, western blot of tissue homogenates revealed that pCD was exclusively detected within tumors (**Fig. 1F**).

### Knockout of *preTA* operon improves the antitumor efficacy of EcNx-pCD/5-FC therapy

After characterizing the EcNx-pCD strain, we proceeded to test its antitumor efficacy. Mice with subcutaneously established MC38 tumors received a single intratumoral (i.t.) injection of either PBS or EcNx-pCD bacteria, followed by i.p. injections of the prodrug 5-FC, as illustrated in **fig. S2**. Unexpectedly, despite validation of the activity of EcNx-pCD/5-FC *in vitro* (**Fig. 1C**), *in vivo* EcNx-pCD/5-FC therapy demonstrated only modest effects on tumor growth suppression (**fig. S2**). Given these contrasting observations, we questioned what factors might be specifically affecting *in vivo* efficacy, with the most obvious difference between *in vitro* and *in vivo* experiments being the length of time that 5-FC was incubated in the presence of EcNx-pCD bacteria. In the *in vivo* setting, both 5-FC and the converted 5-FU were continuously exposed to intratumoral EcNx-pCD bacteria; therefore, we questioned whether bacteria could somehow alter the cytotoxicity of pCD-converted 5-FU chemotherapy. In mammals, 5-FU is detoxified through its metabolic conversion to dihydrofluorouracil (DHFU) by the enzyme dihydropyrimidine dehydrogenase (DPD) (*24*). Patients with partial or complete DPD deficiency are at an increased risk of experiencing severe adverse reactions (*24*). A prior study by Hidese *et al.* found that the *E. coli* DPD encoded by *preT*/*preA* genes (*preTA* operon) is homologous to human DPD and exhibits the ability to catabolize uracil into 5,6-dihydrouracil (*29*). Moreover, subsequent studies have shown that intestinal microbes from CRC patients, as well as clinical isolates of *E.coli* from CRC tumors, carry the functional *preTA* operon and metabolize 5-FU to nontoxic DHFU (*30–32*), thereby impairing 5-FU efficacy. These findings led us to hypothesize that the efficacy of EcNx-pCD/5-FC therapy might be compromised by bacterial detoxification of 5-FU, and that genetic inactivation of the *preTA* operon in EcNx could enhance the stability of 5-FU following intratumoral conversion of prodrug 5-FC by EcNx-pCD (**Fig. 2A**).

**Fig. 2.**
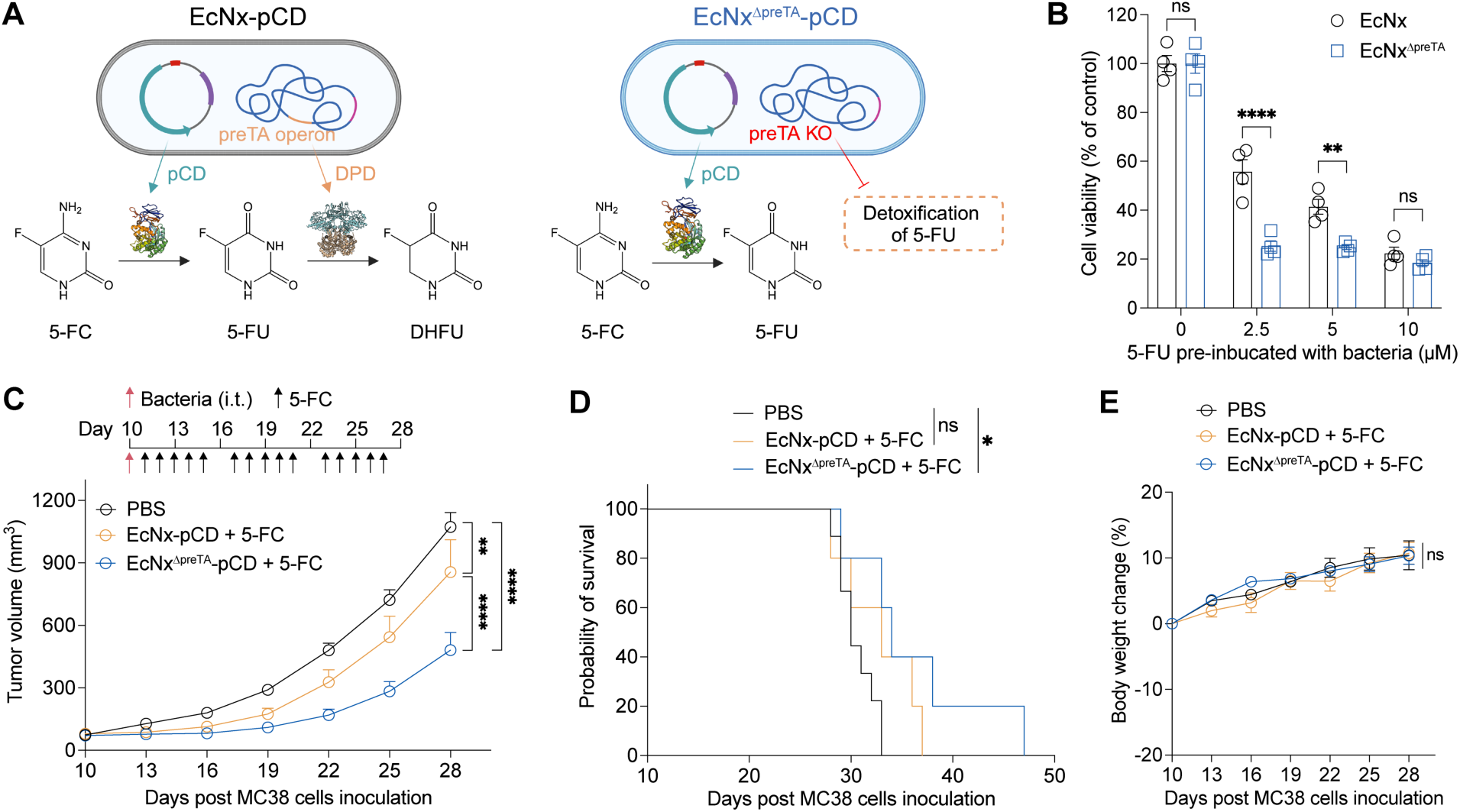
Knockout of preTA operon improves the antitumor efficacy of EcNx-pCD/5-FC. **(A)** The preTA operon of wild-type EcNx was deleted to prevent the detoxification of 5-FU to nontoxic DHFU by preT/preA-encoded dihydropyrimidine dehydrogenase (DPD). The structures of pCD and DPD were modeled by Robetta and AlphaFold3, respectively. **(B)** 5-FU cytotoxicity assay showing the incapability of EcNx^ΔpreTA^ bacteria to detoxify 5-FU. 5-FU was pre-incubated with EcNx or EcNx^ΔpreTA^ bacteria for 5h and then added to MC38 cells. Cell viability was determined using MTT 48h post 5-FU addition. Data are presented as mean ± sem. ns, not significant; ** p = 0.0022; **** p < 0.0001; Two-way ANOVA with Holm-Šídák multiple comparisons test. **(C)** Tumor growth curves and **(D)** survival curves of mice bearing subcutaneous MC38 tumors showing knockout of preTA operon significantly improve the efficacy of EcNx-pCD/5-FC therapy. Data are combined from 3 independent experiments and presented as mean ± sem. (C) ** p = 0.0034; **** p < 0.0001; Two-way ANOVA with Holm-Šídák multiple comparisons test. (D) ns, p = 0.12; * p = 0.02; log-rank (Mantel-Cox) test. **(E)** Body weight change of MC38 tumor-bearing mice in (C) and (D). Data are presented as mean ± sem. ns, not significant; Two-way ANOVA with Holm-Šídák multiple comparisons test.

To obtain the EcNx^ΔpreTA^ strain, we first deleted the coding region of *preTA* operon from wild-type EcN, and then integrated the lysing circuit into the genome of the resultant EcN^ΔpreTA^ strain. After verifying deletion of *preT/preA* genes and integration of the lysing circuit by colony PCR and an *in vitro* lysis, respectively (**fig. S3**), 5-FU detoxification was characterized for both EcNx and EcNx^ΔpreTA^ strains. 5-FU was pre-incubated with EcNx or EcNx^ΔpreTA^ bacteria, and its chemotherapeutic efficacy against MC38 cells was evaluated thereafter. Incubation of 5-FU with EcNx control bacteria showed reduced cytotoxicity compared to incubation with EcNx^ΔpreTA^ bacteria (**Fig. 2B**), indicating that *preTA* knockout effectively abrogates bacteria-mediated 5-FU detoxification. Next, *in vivo* antitumor experiments were conducted to compare the effectiveness of EcNx^ΔpreTA^-pCD/5-FC with that of EcNx-pCD/5-FC (**Fig. 2C and fig. S4**). In line with our prior observation, EcNx-pCD/5-FC demonstrated modest therapeutic efficacy against MC38 tumors, whereas EcNx^ΔpreTA^-pCD/5-FC significantly inhibited tumor growth and correspondingly improved survival compared to PBS control animals (**Fig. 2C, 2D and fig. S4**). Notably, EcNx^ΔpreTA^-pCD/5-FC therapy exhibited no appreciable toxicity, as evidenced by the steady increase in mouse body weight, which was comparable to that of mice in the PBS group, throughout the treatment period (**Fig. 2E**). Altogether, these results indicate that *preTA* knockout blocked the 5-FU detoxification pathway in bacteria, and therefore enhanced CD/5-FC-mediated tumoricidal activity.

### EcNx^ΔpreTA^-pCD/5-FC therapy activates both antitumor effector cells and immunosuppressive cells

Accumulating evidence suggests that chemotherapeutic agents modulate both innate and adaptive antitumor immune responses, in addition to their direct cytotoxicity against tumor cells(*11*). An *in vitro* study by Galetto, *et al*. demonstrated that 5-FU treatment facilitates the uptake of tumor antigens by dendritic cells (DCs) and subsequent antigen cross-presentation to T lymphocytes, resulting in increased release of IFNγ (*33*). Consistently, recent studies using mouse tumor models have shown that 5-FU efficacy requires T cell–mediated antitumor immunity (*13, 34*). Meanwhile, numerous preclinical findings indicate that many chemotherapeutics, including 5-FU, can upregulate PD-L1 expression on tumor cells and tumor-infiltrating myeloid cells, primarily through type I IFN or IFNγ signaling (*12, 35–37*). This PD-L1 upregulation, in turn, impairs the antitumor immune responses elicited by these agents (*12*). An important consideration is that all of these results were obtained with systemically administered 5-FU, and varied depending on the dosage, treatment schedules and tumor models used. For instance, a reduced number of regulatory T cells (Tregs) was observed in the blood of mice treated with a high dosage of 5-FU, whereas low and medium dosages increased the number of circulating Tregs (*38*). Given these discrepancies, we were eager to investigate the immunomodulatory effects of EcNx^ΔpreTA^-pCD/5-FC therapy.

To gain insights into how localized EcNx^ΔpreTA^-pCD/5-FC therapy influences antitumor immune responses, we conducted flow cytometric analysis on tumor-infiltrating immune cells. As illustrated in **Fig. 3A**, mice bearing subcutaneous MC38 tumors received a single intratumoral injection of either EcNx^ΔpreTA^ or EcNx^ΔpreTA^-pCD bacteria, followed by one or two cycles of 5-FC treatment, with each cycle consisting of five consecutive days. Concurrent with the suppression of tumor growth, EcNx^ΔpreTA^-pCD/5-FC therapy led to CD8 T cell activation after one cycle of 5-FC (day 16), as evidenced by a significant increase in IFNγ production compared to PBS and EcNx^ΔpreTA^/5-FC groups (**Fig. 3B**). In addition to CD8 T cells, an increased frequency of IFNγ^+^ cells was also observed in other effector cell populations, including conventional CD4 T (CD4 Tconv) cells, natural killer (NK) cells, and natural killer T (NKT) cells (**Fig. 3C to 3E**). These results suggest that EcNx^ΔpreTA^-pCD/5-FC therapy locally activates antitumor immunity.

**Fig. 3.**
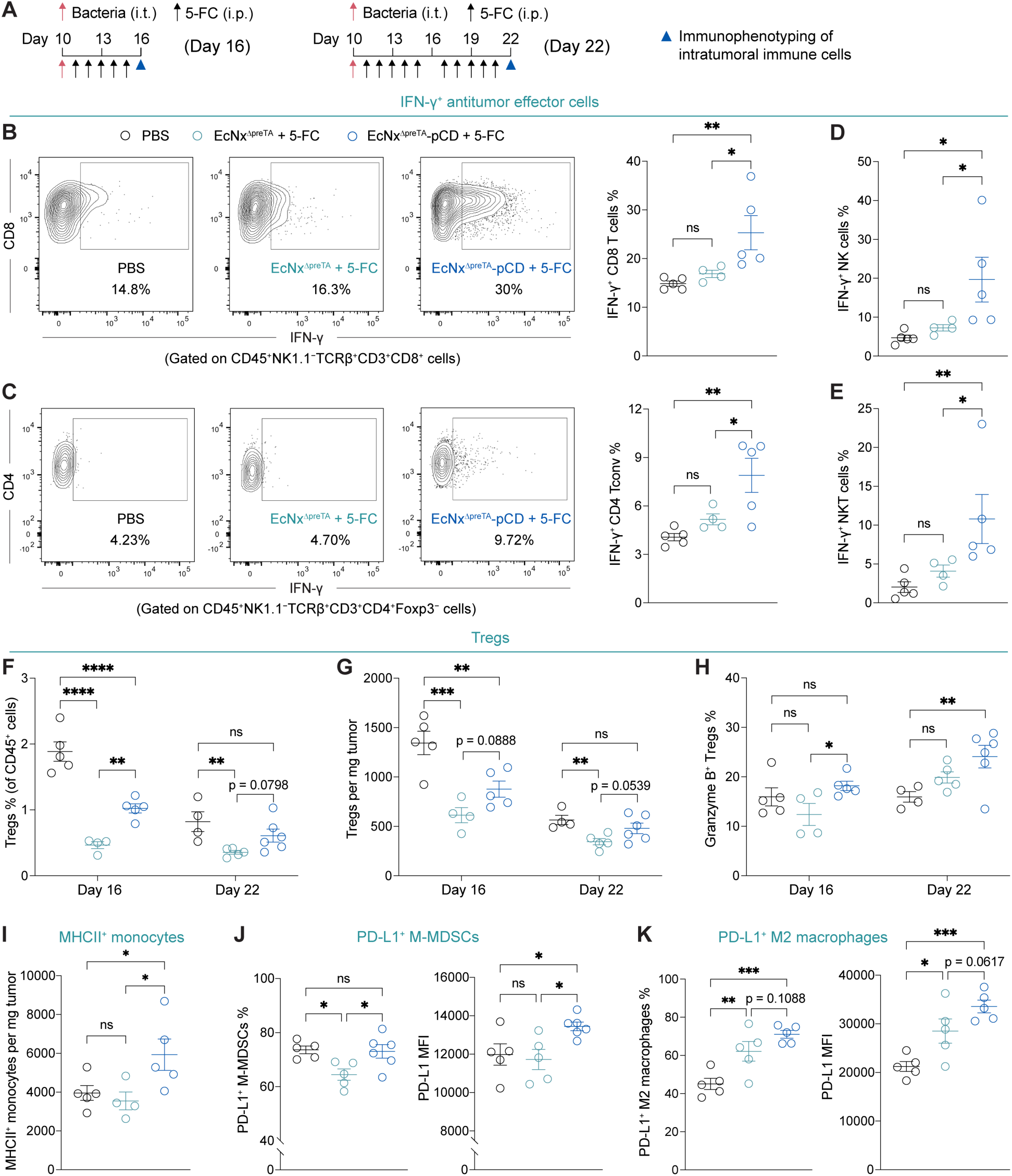
EcNx^ΔpreTA^-pCD/5-FC therapy activates antitumor effector cells and immunosuppressive cells. **(A)** Treatment schedules for immunophenotyping of intratumoral immune cells on day 16 and day 22 post MC38 cells inoculation. C57BL/6NJ mice bearing subcutaneous MC38 tumors received a single intratumoral injection of bacteria, followed by one cycle or two cycles of intraperitoneal 5-FC injection. **(B)** Frequency of IFN-γ^+^ CD8 T (CD45^+^NK1.1^−^CD3^+^TCRβ^+^CD8^+^) cells on Day 16. Representative flow cytometric plots (left) and quantification (right) are shown. **(C)** Frequency of IFN-γ^+^ conventional CD4 T cells (CD4 Tconv, CD45^+^NK1.1^−^CD3^+^TCRβ^+^CD4^+^Foxp3^−^) on Day 16. Representative flow cytometric plots (left) and quantification (right) are shown. **(D)** Frequency of IFN-γ^+^ natural killer cells (NK cells, CD45^+^NK1.1^+^CD3^−^TCRβ^−^) on day 16. **(E)** Frequency of IFN-γ^+^ natural killer T cells (NKT cells, CD45^+^NK1.1^+^CD3^+^TCRβ^+^) on day 16. **(F)** Frequency of Tregs (CD45^+^NK1.1^−^CD3^+^TCRβ^+^CD4^+^Foxp3^+^) of CD45^+^ cells in tumors. **(G)** Number of Tregs per mg tumor. **(H)** Frequency of granzyme B^+^ Tregs. **(I)** Number of MHCII^+^ monocytes (CD45^+^CD11b^+^Ly6G^−^Ly6C^hi^) per mg tumor on day 16. **(J)** Frequency of PD-L1^+^ M-MDSCs (CD45^+^CD11b^+^Ly6G^−^Ly6C^hi^MHCII^−^) and PD-L1 mean fluorescence intensity (MFI) of M-MDSCs on Day 22. **(K)** Frequency of PD-L1^+^ M2 macrophages (CD45^+^CD11b^+^F4/80^+^CD206^+^) and PD-L1 MFI of M2 macrophages on Day 22. (B to I) Data are presented as mean ± sem. ns, not significant; * p < 0.05; ** p < 0.01; *** p < 0.001; **** p < 0.0001; One-way ANOVA with Fisher’s LSD test.

Beyond cytokine production, we next examined whether EcNx^ΔpreTA^-pCD/5-FC therapy can influence the abundance of lymphocytes within tumors. Although no difference in the number of CD8, CD4 Tconv, NK, and NKT cells was observed among three groups (**fig. S5**), tumors injected with EcNx^ΔpreTA^ bacteria showed a reduced number of intratumoral Tregs compared to those treated with PBS on both day 16 and day 22 (**Fig. 3F and 3G**). This observation is consistent with our previous results on EcN-based bacterial therapies (*39*). However, in comparison with EcNx^ΔpreTA^/5-FC, the EcNx^ΔpreTA^-pCD/5-FC group exhibited an elevated frequency of Tregs among CD45^+^ cells, an increased number of Tregs per mg tumor, and a lower CD8/Treg ratio (**Fig. 3F, 3G and fig. S5E**), indicating that this localized chemotherapy facilitates tumor infiltration by Tregs. Furthermore, EcNx^ΔpreTA^-pCD/5-FC therapy resulted in a higher frequency of activated granzyme B-expressing Tregs (**Fig. 3H**), which are known to effectively suppress NK and/or CD8 T cell-mediated antitumor responses (*40, 41*).

We further assessed the impact of EcNx^ΔpreTA^-pCD/5-FC therapy on myeloid cells. Infiltrating monocytes in EcNx^ΔpreTA^-pCD/5-FC-treated tumors showed increased MHCII expression, suggestive of enhanced antigen presentation (**Fig. 3I**). However, the bacterial EPT also augmented the suppressive activity of myeloid cells, as evidenced by the upregulated PD-L1 expression—a classic checkpoint molecule known for its inhibition of cytotoxic T lymphocyte effector function—in both monocytic myeloid-derived suppressor cells (M-MDSCs) and CD206^+^ M2 macrophages (**Fig. 3J and 3K**). Additionally, we found that long-term exposure to bacteria (day 22) elevated PD-L1 expression on intratumoral dendritic cells (DCs) (**fig. S6**). This finding aligns with a previous *in vitro* study demonstrating that stimulation of human CD1c^+^ DCs with *E.coli* increases PD-L1 expression and induces an immunoregulatory phenotype (*42*). Taken together, these results reveal that EcNx^ΔpreTA^-pCD/5-FC therapy activates both antitumor effector cells and suppressor cells.

### Design of EcNx^ΔpreTA^-pCD/PDL1nb/s15 strain for combination chemoimmunotherapy

Considering the effects of EcNx^ΔpreTA^-pCD/5-FC therapy on Tregs and immunosuppressive myeloid cell populations, we reasoned that combining it with immunostimulatory therapy could further enhance its antitumor efficacy. As a member of the IL-2 family cytokines and an alternative to IL-2, IL-15 has been extensively exploited for cancer treatment due to its ability to promote the generation, proliferation, and activity of NK cells and CD8 T cells without supporting tumor-promoting Tregs (*43*). More importantly, IL-15 has been reported to render conventional T lymphocytes resistant to the suppressive effects of Tregs, thereby sustaining their proliferation and IFNγ production (*44, 45*). Inspired by these findings and our observations of Treg activation and PD-L1 upregulation upon EcNx^ΔpreTA^-pCD/5-FC treatment, we sought to co-express an IL-15 superagonist (IL-15Rα_sushi_-IL-15 fusion protein, referred to as “s15”) (*46*) and a PD-L1 nanobody (PDL1nb) (*20*) alongside pCD on a single plasmid (**Fig. 4A**). We hypothesized that continuous production of s15 and PDL1nb by bacteria within the TME could overcome the immunosuppressive effects of Tregs and block PD-L1 inhibitory signaling, respectively, enhancing the therapeutic effect of EcNx-pCD/5-FC.

**Fig. 4.**
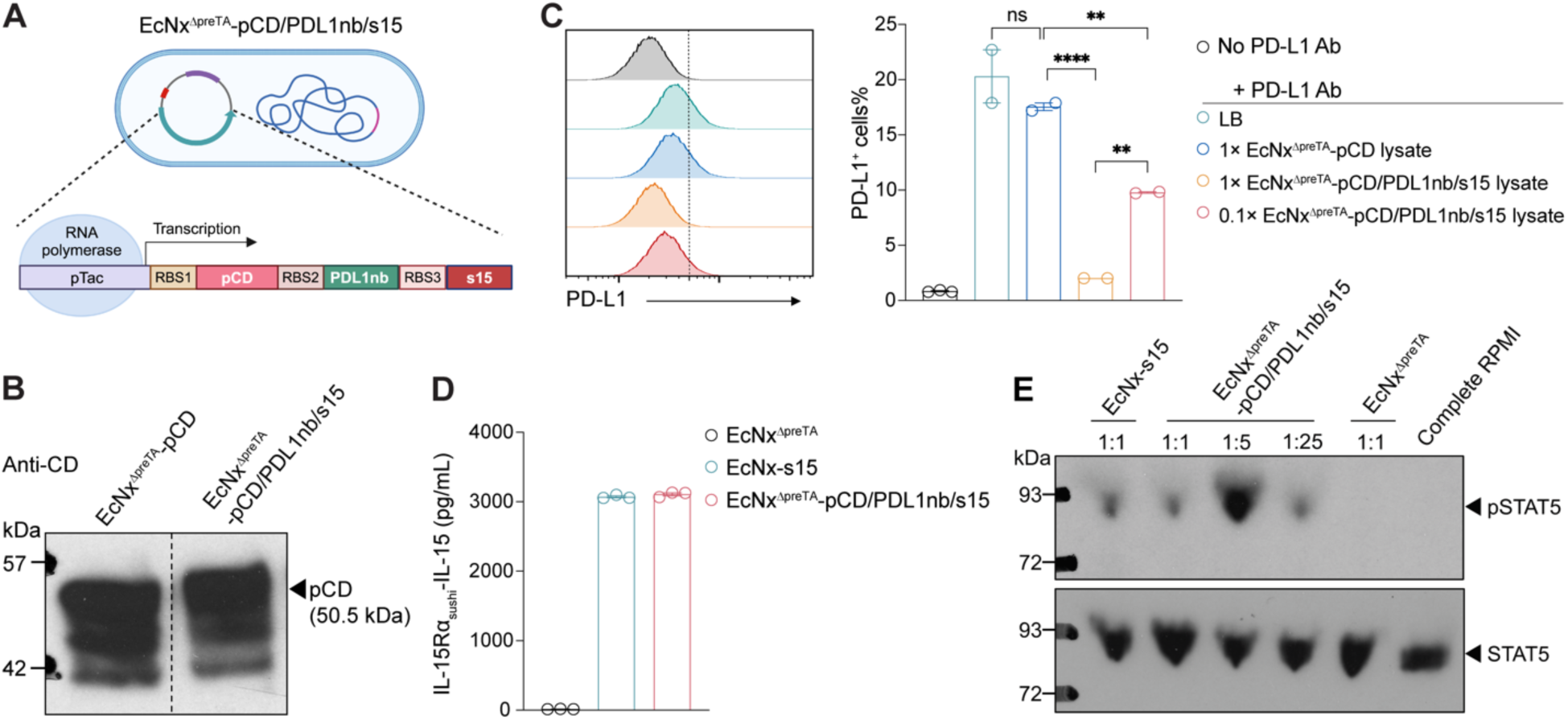
Characterization of EcNx^ΔpreTA^-pCD/PDL1nb/s15 strain. **(A)** Schematic illustration showing the co-expression of pCD, PD-L1 nanobody (PDL1nb) and IL-15Rα_sushi_-IL-15 (s15) fusion protein in EcNx^ΔpreTA^ bacteria. **(B)** Western blot image showing the expression of pCD by EcNx^ΔpreTA^-pCD and EcNx^ΔpreTA^-pCD/PDL1nb/s15 strains. The dashed black line indicates the cropping to remove a non-relevant lane. **(C)** Merged flow cytometric histogram (left) and percentage of PD-L1^+^ MC38 cells (right) showing the production and activity of PDL1nb. MC38 cells were pre-incubated with the lysates of indicated strains for 20 min and then stained with APC-labeled PD-L1 antibody for flow cytometric analysis. LB, lysogeny broth. **(D)** ELISA quantification of IL-15Rα_sushi_-IL-15 concentration in the lysates of EcNx^ΔpreTA^, EcNx-s15 and EcNx^ΔpreTA^-pCD/PDL1nb/s15 bacteria. **(E)** Western blot detection of phosphorylated STAT5 (pSTAT5) and STAT5. Freshly isolated splenocytes were stimulated with indicated bacteria lysates for 30 min and then collected for western blot. (C and D) Data are presented as mean ± sem. (C) ns, not significant; ** p < 0.01; **** p < 0.0001; One-way ANOVA with Fisher’s LSD test.

We first validated the bacterial co-expression of pCD, PDL1nb, and s15 by transforming the Axe/Txe-stabilized p246-pCD/PDL1nb/s15 plasmid into both wild-type EcNx and EcNx^ΔpreTA^ bacteria. As shown by western blot analysis, EcNx/EcNx^ΔpreTA^-pCD/PDL1nb/s15 bacteria produced comparable levels of pCD to EcNx/EcNx^ΔpreTA^-pCD strains (**Fig. 4B and fig. S7A**), suggesting that the co-expression of PDL1nb and s15 has minimal effect on pCD production. To verify the production and bioactivity of PDL1nb, we performed a cell surface PD-L1 binding assay using bacterial lysates and an APC-conjugated PD-L1 mAb (clone 10F.9G2), which recognizes the same epitope as PDL1nb (*20*). MC38 cells were incubated with the lysates from different bacterial strains for 20 minutes, then stained with the fluorescent PD-L1 mAb for flow cytometric analysis. While cells pre-incubated with LB medium and EcNx/EcNx^ΔpreTA^-pCD lysates displayed PD-L1 mAb fluorescence, EcNx/EcNx^ΔpreTA^-pCD/PDL1nb/s15 lysates significantly blocked mAb binding to PD-L1 on MC38 cells, as demonstrated by a dose-dependent decrease in mAb fluorescence with increasing lysate concentrations (**Fig. 4C and fig. S7B**). These results indicate the successful production of functional PDL1nb by both EcNx- and EcNx^ΔpreTA^-pCD/PDL1nb/s15 bacteria. ELISA quantification of s15 in bacterial lysates further confirmed that this co-expression plasmid allowed s15 production at a level comparable to the strain expressing s15 alone (EcNx-s15) (**Fig. 4D and fig. S7C**). Moreover, STAT5 phosphorylation was observed in mouse splenocytes stimulated with EcNx-s15 and EcNx^ΔpreTA^-pCD/PDL1nb/s15 lysates but not with EcNx^ΔpreTA^ lysate, indicating the bioactivity of s15 (**Fig. 4E**). Collectively, these data indicate we have successfully engineered a versatile strain capable of simultaneously producing the prodrug-catalyzing enzyme cytosine deaminase, PD-L1 blocking nanobody, and s15 cytokine, presenting a promising platform for chemoimmunotherapy combination.

### Bacterial combination chemoimmunotherapy boosts both adaptive and innate antitumor immunity

We next proceeded to assess the antitumor efficacy of the chemoimmunotherapy combination strategy. Initial experiments on MC38 tumor growth were conducted using the wild-type EcNx-based strains. Although the EcNx-pCD/5-FC therapy exhibited marginal suppression of MC38 tumors as previously observed (**fig. S2 and S7D**), the addition of PDL1nb and s15 considerably improved its effectiveness (**fig. S7D and S7F**). Likewise, this augmentation was observed in the experiments using *preTA* knockout strains. A single i.t. injection of EcNx^ΔpreTA^-pCD/PDL1nb/s15 bacteria, together with subsequent i.p. injections of 5-FC, resulted in significantly enhanced tumor control compared to both EcNx^ΔpreTA^-pCD/5-FC treatment and EcNx^ΔpreTA^-pCD/PDL1nb/s15 bacteria alone (**Fig. 5A and fig. S8A**). This indicates that cytotoxic 5-FU, converted from 5-FC prodrug, synergizes with PDL1nb and s15 to induce effective antitumor responses. Remarkably, 3 out of 7 tumors treated with EcNx^ΔpreTA^-pCD/PDL1nb/s15 bacteria plus 5-FC showed a complete regression (**Fig. 5A**), whereas combination treatment with wild-type EcNx failed to eradicate any established tumors (**fig. S7F**), again highlighting the necessity of *preTA* knockout. To evaluate the therapeutic benefit of the combination approach in a more translational context, we treated tumor-bearing mice with a single dose of bacteria via tail vein injection and subsequently administered 5-FC, as illustrated in **Fig. 5B**. Consistent with the results of i.t. injection, intravenously-delivered EcNx^ΔpreTA^-pCD/PDL1nb/s15 bacteria markedly inhibited tumor growth in the presence of 5-FC (**Fig. 5B and fig. S9**). Meanwhile, EcNx^ΔpreTA^-pCD/PDL1nb/s15 bacteria alone, or EcNx^ΔpreTA^-pCD/5-FC, demonstrated only moderate efficacy (**Fig. 5B**). Most notably, for both routes of bacterial administration, no obvious decrease in mouse body weight was observed during the experimental period, suggesting that this tumor-localized chemoimmunotherapy combination was well-tolerated (**Fig. 5C, fig. S7E and S8B**).

**Fig. 5.**
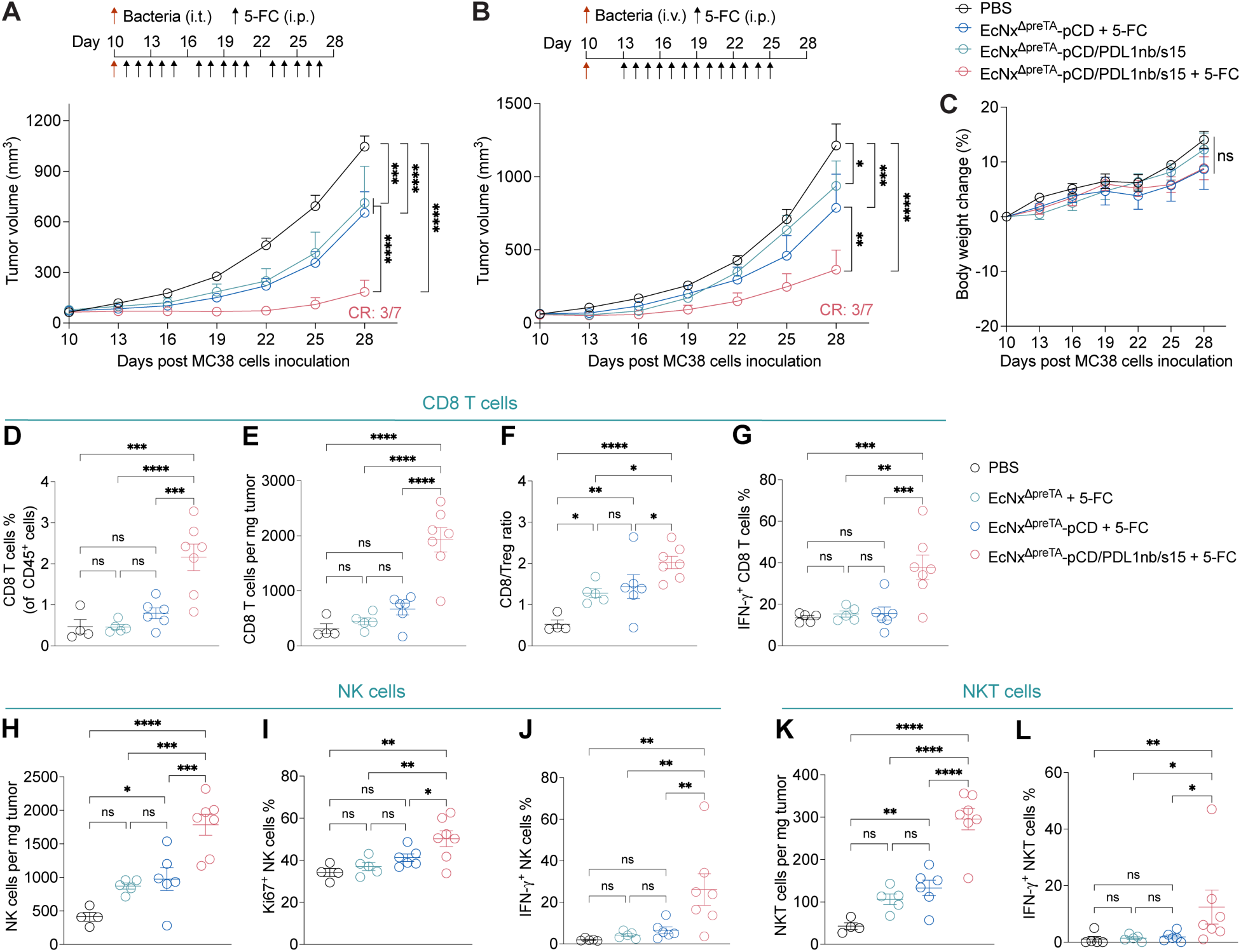
Bacterial combination chemoimmunotherapy boosts both adaptive and innate antitumor immunity. **(A and B)** MC38 tumor growth curves. C57BL/6NJ mice received either (A) intratumoral or (B) intravenous injection of indicated bacteria (1 × 10^6^ CFU/tumor and 2 × 10^7^ CFU/mouse, respectively) on day 10 post subcutaneous inoculation of MC38 cells, followed by intraperitoneal injections of 5-FC (500 mg/kg) as illustrated (black arrows). **(C)** Body weight changes of mice in (B). (A to C) Data are presented as mean ± sem. ns, not significant; * p < 0.05; *** p < 0.001; **** p < 0.0001; Two-way ANOVA with Holm-Šídák multiple comparisons test. **(D to L)** Mice bearing MC38 tumors received intratumoral injection of either EcNx^ΔpreTA^, EcNx^ΔpreTA^-pCD, or EcNx^ΔpreTA^-pCD/PDL1nb/s15 bacteria (1 × 10^6^ CFU/tumor) and intraperitoneal injections of 5-FC (500 mg/kg) as illustrated in (A). Tumors were collected on day 22 for flow cytometric analysis of tumor-infiltrating lymphocytes. **(D)** Frequency of CD8 T cells of CD45^+^ cells. **(E)** Number of CD8 T cells per mg tumor. **(F)** Ratio of intratumoral CD8 T cells to Tregs. **(G)** Frequency of IFN-γ^+^ CD8 T cells. **(H)** Number of NK cells per mg tumor. Frequency of **(I)** Ki67^+^ and **(J)** IFN-γ^+^ NK cells. **(K)** Number of NKT cells per mg tumor. **(L)** Frequency of IFN-γ^+^ NKT cells. (D to L) Data are presented as mean ± sem. ns, not significant; * p < 0.05; ** p < 0.01; *** p < 0.001; **** p < 0.0001; One-way ANOVA with Fisher’s LSD test.

To elucidate how the addition of PDL1nb and s15 contributes to antitumor immunity, we treated MC38 tumors as shown in Figure 5A and harvested tumors on day 22 for flow cytometric analysis of infiltrating immune cells (**fig. S10**). EcNx^ΔpreTA^-pCD/PDL1nb/s15 bacteria plus 5-FC treatment led to a substantial increase in the frequency of cytotoxic CD8 T cells among CD45^+^ cells and their density per mg tumor, whereas EcNx^ΔpreTA^-pCD/5-FC therapy elicited only a slight but non-statistically significant trend when compared to PBS and EcNx^ΔpreTA^ bacteria (**Fig. 5D and 5E**). In accordance with the expansion of CD8 T cells, tumors treated with the combination therapy displayed the highest CD8/Tregs ratio among all four groups (**Fig. 5F**). Interestingly, although we observed the activation of CD8 T cells on day 16 following EcNx^ΔpreTA^-pCD/5-FC treatment (**Fig. 3B**), by day 22, these cells showed no increase in IFNγ production (**Fig. 5G**), implying that prolonged EcNx^ΔpreTA^-pCD/5-FC treatment might compromise CD8 T cell functionality. This cytokine production deficit was fully restored upon combining EcNx^ΔpreTA^-pCD/5-FC with PDL1nb and s15 (**Fig. 5G**). Moreover, the combination therapy considerably increased the number of NK and NKT cells, while driving their production of IFNγ, and promoting NK cell proliferation concurrently (**Fig. 5H to 5L**). In addition to stimulating effector cells, the chemoimmunotherapy combination reduced the frequency of M-MDSCs (**fig. S11**), a key immunosuppressive cell population that inhibits antitumor immune responses (*47*).

Given the central role of antigen presentation in initiating and maintaining antitumor T cell immunity (*48*), we then investigated the status of antigen-presenting cells (APCs) after treatment with EcNx^ΔpreTA^-pCD/PDL1nb/s15 bacteria plus 5-FC. We found that in the presence of PDL1nb and s15, type 2 conventional dendritic cells (cDC2), monocyte-derived dendritic cells (moDCs) and macrophages upregulated major histocompatibility complex class II (MHCII) as well as the costimulatory molecules CD80 and CD86 (**Fig. 6A to 6C and fig. S12**). For type 1 conventional dendritic cells (cDC1), while bacterial stimulation alone was able to induce CD86 expression, increases in MHCII and CD80 MFIs were observed only in the EcNx^ΔpreTA^-pCD/PDL1nb/s15 plus 5-FC group (**Fig. 6A to 6C**). Furthermore, the combination therapy skewed macrophages toward the M1 phenotype, as indicated by the increased frequency and number of MHCII^+^CD80^+^ macrophages within tumors (**Fig. 6D and 6E**). M1 cells in the combination group also displayed higher MHCII MFI than those in other groups (**Fig. 6F**). Overall, these results demonstrate that the addition of PDL1nb and s15 promotes dendritic cell activation and macrophage M1 polarization, thereby enhancing their antigen-presenting capability.

**Fig. 6.**
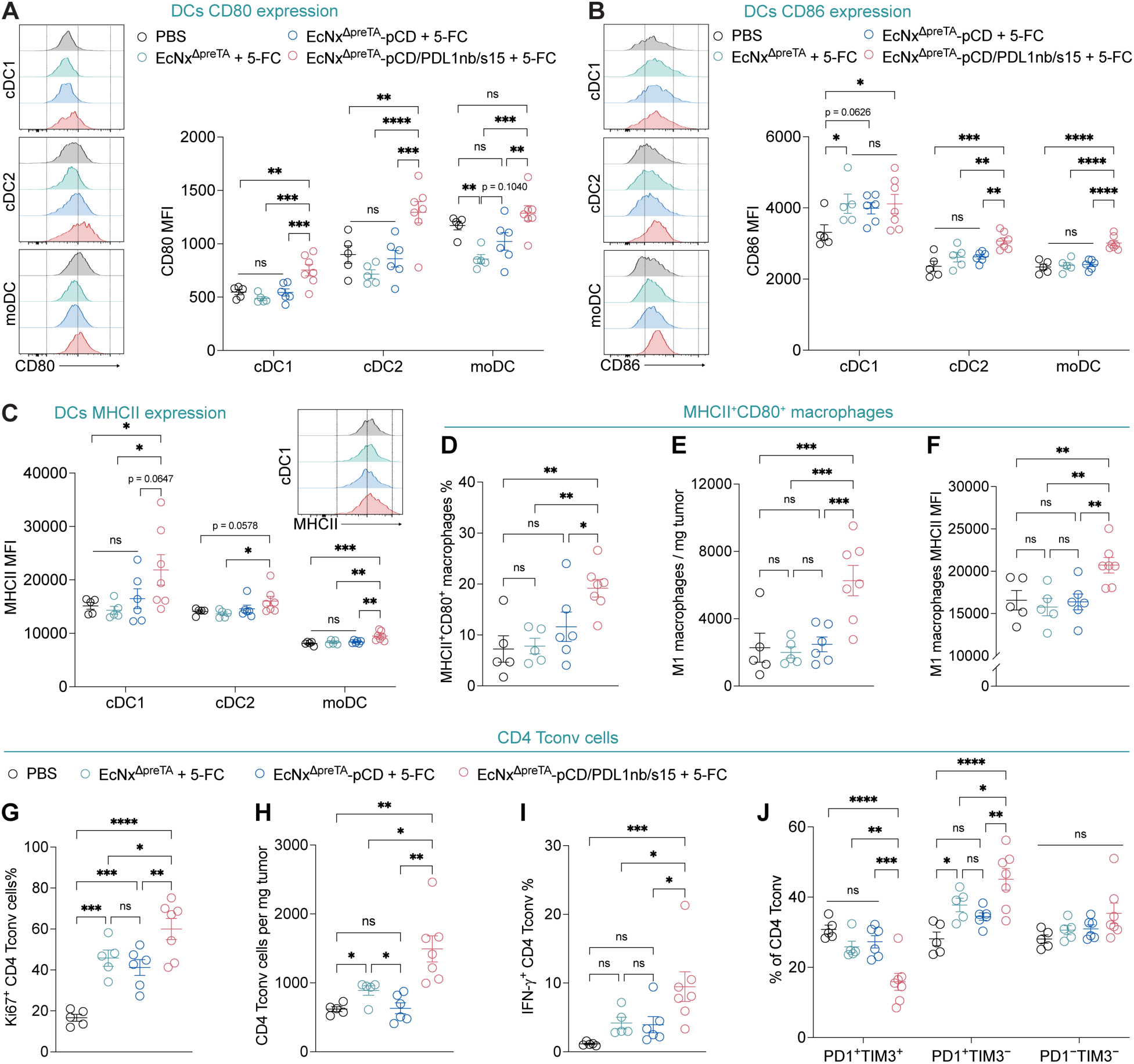
Bacterial combination chemoimmunotherapy activates antigen-presenting cells and reprograms conventional CD4 T cells. C57BL/6NJ mice bearing subcutaneous MC38 tumors received intratumoral injection of either EcNx^ΔpreTA^, EcNx^ΔpreTA^-pCD, or EcNx^ΔpreTA^-pCD/PDL1nb/s15 bacteria (1 × 10^6^ CFU/tumor) and intraperitoneal injections of 5-FC (500 mg/kg) as illustrated in figure 5A. Tumors were collected on day 22 post-cancer cell inoculation for flow cytometric analysis of intratumoral immune cells. Representative histograms (left) and MFIs (right) showing the expression of **(A)** CD80 and **(B)** CD86 on dendritic cell populations (cDC1, type 1 conventional dendritic cells, CD45^+^Ly6G^−^Ly6C^−^F4/80^−^CD11c^+^MHCII^+^CD103^+^; cDC2, type 2 conventional dendritic cells, CD45^+^Ly6G^−^Ly6C^−^F4/80^−^CD11c^+^MHCII^+^CD11b^+^CD103^−^; moDCs, monocytes-derived dendritic cells, CD45^+^Ly6G^−^Ly6C^+^CD11c^+^MHCII^+^). **(C)** MHCII MFI of different dendritic cell populations. **(D)** Frequency of MHCII^+^CD80^+^ M1 macrophages. **(E)** Number of M1 macrophages per mg tumor tissue. **(F)** MHCII MFI of M1 macrophages. **(G)** Frequency of Ki67^+^ CD4 Tconv cells. **(H)** Number of CD4 Tconv cells per mg tumor tissue. **(I)** Frequency of IFN-γ^+^ CD4 Tconv cells. **(J)** Frequency of PD-1^+^TIM-3^+^, PD-1^+^TIM-3^−^ and PD-1^−^TIM-3^−^ cells of CD4 Tconv cells. (A to J) Data are presented as mean ± sem. ns, not significant; * p < 0.05; ** p < 0.01; *** p < 0.001; **** p < 0.0001; One-way ANOVA with Fisher’s LSD test.

Growing evidence indicates that CD4 Tconv cells play a critical role in orchestrating the crosstalk between innate and adaptive immune cells for optimal antitumor immunity (*49, 50*). Activated APCs can prime CD4 T cells, which in turn amplify APC activation through the upregulation of CD40 ligand (CD40L) (*50*). This process “licenses” APCs to support clonal expansion and effector/memory differentiation of CD8 T cells (*50, 51*). Additionally, CD4 Tconv cells secret cytokines, such as IFNγ, to enhance the proliferation and cytotoxicity of CD8 T cells, NK cells and NKT cells (*49*). Based on the activation of APCs and antitumor effector cells observed with the combination therapy, we hypothesized that bacteria-produced PDL1nb and s15 might have a profound impact on conventional CD4 T cells. Indeed, although the bacterial vector alone was able to boost the proliferation of CD4 Tconv cells, presumably due to its induction of CD86 on cDC1 (**Fig. 6B**), the chemoimmunotherapy combination resulted in a response of significantly higher magnitude (**Fig. 6G**). Intriguingly, while tumors treated with EcNx^ΔpreTA^ bacteria plus 5-FC showed a corresponding increase in the number of CD4 Tconv cells compared to the PBS control, no such increase was observed with the EcNx^ΔpreTA^-pCD plus 5-FC treatment (**Fig. 6H**). This implies that the persistence of 5-FU within TME might promote CD4 Tconv cell death. Notably, combining with PDL1nb and s15 completely rescued the expansion of CD4 Tconv cells while also enhancing their IFNγ production (**Fig. 6H and 6I**). CD4 T cell exhaustion has been reported to negatively influence CD4 T cell proliferation and cytokine production, B cell help, and CD8 effector functions, thus hindering antitumor immunity (*52*). This exhaustion is commonly accompanied by the upregulation of PD-1 and T cell immunoglobulin and mucin domain 3 (TIM-3) (*52*). Our immune phenotyping analysis revealed that PD-1^+^TIM-3^+^, PD-1^+^TIM-3^−^ and PD-1^−^TIM-3^−^ populations of CD4 Tconv cells were present at roughly equal frequency in the tumors treated with PBS, and treatment with EcNx^ΔpreTA^ bacteria plus 5-FC or EcNx^ΔpreTA^-pCD plus 5-FC did not dramatically alter their proportions (**Fig. 6J**). In contrast, EcNx^ΔpreTA^-pCD/PDL1nb/s15 plus 5-FC significantly reduced the frequency of PD-1^+^TIM-3^+^ CD4 Tconv cells and at the same time increased the proportion of PD-1^+^TIM-3^−^ cells (**Fig. 6J**), suggesting a reversal of CD4 Tconv cell exhaustion. Altogether, these results indicate that the addition of PDL1nb and s15 promotes proliferation and effector function of CD4 Tconv cells and skews them away from an exhausted phenotype.

## DISCUSSION

In this report, we have engineered the probiotic *E.coli* Nissle 1917 to couple enzyme/prodrug strategy with tumor-targeted delivery of immunotherapeutics, establishing a microbial platform for localized cancer chemoimmunotherapy.

Enzyme/prodrug systems have been investigated for decades as a means to enhance the efficacy and safety of conventional chemotherapies, though with limited success in the clinic (*53*). One key challenge is the tumor delivery of prodrug-activating enzymes, which is typically achieved through gene delivery or conjugation to tumor-targeting antibodies (*2, 3*). However, virus-based vectors face safety concerns and host immune responses against viral particles further limit durable gene expression. Similarly, enzyme-antibody conjugates often require repeated administration due to their instability and fast degradation. Furthermore, none of these approaches demonstrate both efficient tumor penetration and satisfactory tumor selectivity. Unlike traditional vectors, the intrinsic tumor-colonizing ability of living EcN bacteria makes them an ideal vehicle to deliver candidate enzymes for tumor-localized EPT (*18, 54*). We show here that the enzyme CD can be exclusively produced within tumor tissues following a single dose of CD-expressing bacteria (**Fig. 1D to 1F**). Upon systemic administration of the prodrug 5-FC, this bacteria-based EPT effectively controls MC38 tumor growth without causing toxicity in mice (**Fig. 2E**), highlighting its translational potential. Notably, a recent phase I study of SYNB1891, an engineered EcN strain expressing STING agonist suggested that repeat intratumoral injection of EcN is safe and well-tolerated in patients with advanced malignancies (*55*).

A successful EPT depends not only on tumor delivery of enzymes but also on efficient prodrug conversion, which is largely determined by the catalytic activity of the enzyme and its selectivity for prodrug molecules. A past trial of the engineered *Salmonella* strain VNP20009 expressing *E. coli* CD demonstrated bacterial tumor colonization in 2 out of 3 refractory cancer patients with acceptable safety profiles (*56*). However, the clinical responses were modest, presumably due to the low conversion efficiency of 5-FC to 5-FU (*56*). To address this issue, we employ a previously reported *E. coli* CD mutant in our study, which displays a significant substrate preference toward 5-FC over cytosine (*28*). For eventual clinical applications, directed evolution may be applied to further improve enzyme efficiency, reducing the required prodrug dose while enhancing the efficacy. Additionally, although many different bacterial strains have been explored for EPT, their intrinsic metabolic effect on the activated drugs, a potentially critical factor influencing EPT efficacy, have not received sufficient attention. Recent studies on the metabolism of anti-cancer drugs by gut or intratumoral bacteria reveal that the *E. coli* enzyme DPD detoxifies 5-FU to DHFU, thereby reducing 5-FU bioavailability (*31, 32*). Inspired by these findings, we deleted the DPD-encoding *preTA* operon in the EcNx strain to abolish 5-FU detoxification. Our results demonstrated that *preTA* knockout significantly improves the *in vivo* efficacy of EcNx-pCD/5-FC therapy (**Fig. 2C and 2D**). This suggest that, in addition to enzyme kinetics, metabolism of the activated drug by bacterial vectors should be considered when designing a bacterial EPT.

Another notable feature of our platform is the rational combination of EPT with immunotherapy. Despite its reduced systemic toxicity and enhanced efficacy, EPT still faces the challenge of drug resistance, a common issue in conventional chemotherapy. Unlike chemotherapeutics, immunotherapy agents harness the host’s immune system to recognize and destroy cancer cells. Nevertheless, their efficacy largely depends on tumor immunogenicity and pre-existing antitumor immunity. The distinct mechanisms of EPT and immunotherapy make their combination an attractive strategy to address the limitations of each. Although certain chemotherapeutics have demonstrated synergy with immune checkpoint blockades, it is important to note that the delivery method of chemotherapy drugs significantly influences their effects on the host’s immune system. Systemic chemotherapy has been shown to compromise peripheral immune integrity and hinder the effectiveness of PD-1 blockades (*36, 57*), whereas locally administered chemotherapy drugs enhance antitumor immune responses (*57*). These findings highlight the potential advantages of EPT in stimulating local immune responses, while preserving intact peripheral immunity. Our study demonstrates that EcNx^ΔpreTA^-pCD/5-FC therapy activates antitumor T cells, NK cells, and NKT cells within the tumor microenvironment (**Fig. 3A to 3E**). However, a thorough examination of other immune cell types revealed some negative immunologic effects, including the activation of Tregs and upregulation of PD-L1 on suppressor cells (**Fig. 3F to 3H, 3J and 3K**). Building on these findings, we further refined the EcNx^ΔpreTA^-pCD/5-FC therapy by co-producing an IL-15 superagonist and a PD-L1 blocking nanobody in the bacteria. This combination facilitated the activation of both innate and adaptive immune cells within tumors, leading to enhanced antitumor efficacy in syngeneic mouse models (**Fig. 5 and 6**). The emphasis on local delivery and rational combination design offers a promising direction for future studies to optimize chemoimmunotherapy approaches.

Taken together, our results highlight the potential of living bacteria as a potent platform to integrate enzyme/prodrug therapy and immunotherapeutics, including cytokines and immune checkpoint blockade, for precision cancer treatment. By addressing the challenges of tumor specificity and immunosuppression, this probiotic platform provides a comprehensive solution to limitations faced by existing therapies. Additionally, its modularity allows for seamless incorporation of diverse enzyme/prodrug candidates and immune-activating molecules. With a deeper understanding of tumor immunology and emerging biomarkers, this system should help advance cancer chemoimmunotherapy by offering a highly adaptable, targeted, and synergistic approach to overcoming tumor heterogeneity, toxicities, and treatment resistance across a variety of cancer types.

## MATERIALS AND METHODS

### Study Design

The objective of this study was to develop a probiotic platform for tumor-localized combination chemoimmunotherapy. To achieve this, cytosine deaminase was expressed in bacteria to locally convert the prodrug 5-FC into the cytotoxic agent 5-FU within tumors. Additionally, an IL-15 superagonist and a PD-L1-blocking nanobody were co-expressed to enhance antitumor immune responses. The *preTA* operon was deleted from the bacterial genome using the lambda Red recombination system to prevent 5-FU detoxification. The bioactivity of cytosine deaminase and the PD-L1-blocking nanobody was assessed *in vitro* using MC38 cells via a cell viability assay and cell surface PD-L1 staining, respectively. The functionality of the IL-15 superagonist was evaluated in splenocytes through pSTAT5 detection by western blot. The *in vivo* antitumor efficacy of the probiotic platform was tested in a subcutaneous MC38 tumor model using both intratumoral and intravenous delivery. Mice were grouped based on comparable tumor volumes at the start of treatment to ensure uniform baseline tumor sizes. Tumor growth was monitored via caliper measurements, and mouse body weight was recorded as an indicator of overall health. Analysis of intratumoral immune cells was performed using flow cytometry. Investigators were not blinded to group assignments for *in vitro* and *in vivo* studies, as this information was essential for conducting experiments. Sample sizes and statistical analyses were determined based on field standards and our prior experience with bacterial and mammalian cell studies, as well as animal trials (*19–23, 39*). The sample size for each experimental group is explicitly stated in the figures, figure legends, or data descriptions.

### Mice

7-week-old female C57BL/6NJ (strain 005304) mice were purchased from the Jackson Laboratory, and housed in a temperature-controlled room on a 12:12 h light/dark cycle in the animal house maintained by Institute of Comparative Medicine (ICM), Columbia University. All animal procedures were conducted with the approval of the Institutional Animal Care and Use Committee of Columbia University (protocol AC-AABT2656).

### Cell lines

MC38 cells were maintained in Dulbecco’s Modified Eagle Medium (DMEM) supplemented with 10% FBS (Corning), 100 U/mL Penicillin-Streptomycin (Gibco), 1× MEM nonessential amino acids (Gibco), 1× GlutaMAX (Gibco), 10 mM HEPES (Gibco) at 37℃ in a humidified incubator containing 5% CO_2_.

### EcNx^ΔpreTA^ strain

Genetic knockout of *preTA* operon was conducted using the lambda red recombination system (*58*). In summary, *E.coli* Nissle 1917 was transformed with the pKD46 plasmid. The transformants were cultured at 30°C in LB media containing ampicillin and L-arabinose, followed by preparation of electrocompetent cells. A chloramphenicol resistance cassette with homologous arms flanking the target gene was generated by PCR amplification from pKD3. Electroporation was performed using 100 μL of competent cells and 100-200 ng of the amplified DNA. After a 2-hour recovery in SOC media, cells were plated on LB agar supplemented with chloramphenicol and incubated overnight at 37°C. The gene deletion was confirmed using colony PCR. A synchronized lysing circuit was then integrated into the genome of EcN^ΔpreTA^ as previously described (*20*) to obtain the EcNx^ΔpreTA^ strain.

### Plasmids

Gene fragments of PIGF2_123-144_-tagged bacterial cytosine deaminase mutant 1525 (V152A, F316C and D317G) (pCD)(*19, 28*), PD-L1 nanobody (PDL1nb, RCSB PDB: 5DXW) and human IL-15Rα_sushi_-IL-15 fusion protein (s15) (*46*) were synthesized (Twist Bioscience) and cloned into plasmid p246 (*20*) via Gibson Assembly (NEB). The resultant p246-pCD, p246-pCD/PDL1nb, and p246-pCD/PDL1nb/s15 plasmids were transformed into EcNx or EcNx^ΔpreTA^ bacteria via electroporation. Empty EcNx and EcNx^ΔpreTA^ bacteria were grown in LB broth with erythromycin (100 μg/mL). Strains with a therapeutic plasmid (p246) were grown in LB broth with kanamycin (50 μg/mL).

### 5-FU catabolism

10 mM 5-FU was incubated with 4 × 10^9^ CFU/mL EcNx or EcNx^ΔpreTA^ bacteria in 1 mL serum-free DMEM at 37 ℃ for 5 hours. After pelleting bacteria down, the supernatants were diluted to 10, 5, and 2.5 µM in complete DMEM and added into MC38 cells in a 96-well plate. MC38 cells were incubate with the EcNx/EcNx^ΔpreTA^-conditioned 5-FU for 48 hours, followed by cell viability determination using MTT reagent (Sigma Aldrich, Cat. No. M2128).

### Characterization of pCD

To examine the expression of pCD, lysates of EcNx/EcNx^ΔpreTA^ and pCD-expressing strains were prepared via boil of cell pellets in 1× SDS loading dye for Western Blot (WB). Briefly, lysates in optimal volume were loaded to a Bolt™ 4-12% Bis-Tris Plus gel for SDS-PAGE, and the proteins were transferred to a PVDF membrane. The recombinant pCD was detected using anti-bacterial cytosine deaminase antibody (GeneTex, Cat. No. GTX124166).

To assay for the enzymatic activity of pCD, EcNx and EcNx-pCD bacteria were washed with cold PBS, matched at optical density at 600 nm (OD_600_), and resuspended in 1 mL DMEM containing 2 mM 5-FC, followed by a co-incubation at 37℃ for 30 min. Bacteria suspensions were then centrifuged at 14,000 rpm for 10 min, and the resultant supernatants were diluted and added into MC38 cells in a 96-well plate. After 48 h incubation at 37℃, cell viability was determined using MTT reagent (Sigma Aldrich, Cat. No. M2128).

### Bioactivity of PDL1nb

To verify the PD-L1 binding ability of PD-L1nb produced by EcNx/EcNx^ΔpreTA^ bacteria, MC38 cells (3 × 10^5^ cells/well) were incubated with 100 uL/well of bacterial lysates of relevant strains in a 96-well V-bottom plate at 4℃ for 30 min, followed by staining of MC38 cells using APC-conjugated monoclonal PD-L1 antibody (clone 10F.9G2, BioLegend). Subsequent flow cytometric analyses were performed on a BD LSRFortessa cell analyzer, and the data were analyzed using FlowJo software.

### Characterization of IL-15Rα_sushi_-IL-15 fusion protein

The bacterial production of s15 was confirmed by an IL-15 enzyme-linked immunosorbent assay (ELISA). 1 mL overnight culture of EcNx^ΔpreTA^, EcNx-s15 or EcNx^ΔpreTA^-pCD/PDL1nb/s15 bacteria was inoculated into 50 mL LB with 50 µg/mL kanamycin and then subcultured at 37℃ for 3 h. Bacteria were washed twice with cold PBS, matched at OD_600_, and lysed via sonication in 3 mL complete RPMI-1640. Lysate was centrifuged at 14,000 rpm for 20 min at 4℃ to remove debris, and the supernatant was entered into an ELISA detection of IL-15 (BioLegend ELISA MAX™ Deluxe Set Human IL-15 kit, Cat # 435104).

To test the bioactivity of s15, 1 mL bacterial lysates of EcNx^ΔpreTA^, EcNx-s15 or EcNx^ΔpreTA^-pCD/PDL1nb/s15 strains were prepared as described above, and incubated with 15 million splenocytes at 37℃ for 30 min. Cells were then collected by centrifugation (450*g*, 5 min), and boiled in 1× SDS loading dye for WB detection of pSTAT5 (Cell Signaling, Phospho-Stat5 (Tyr694) Antibody #9351).

### Animal experiments

7-week-old mice purchased from Jackson lab were allowed to acclimate for a week, and then subcutaneously injected with 100 uL MC38 cell suspension (5 × 10^6^ cells/mL in serum-free DMEM). The length and width of tumors were measured using a digital caliper, and the tumor volume was calculated based on the following formula: tumor volume = (length × width^2^)/2. For tumor treatment with bacteria, bacteria were cultured from glycerol stocks overnight in 5 mL LB with appropriate antibiotics at room temperature, and then subcultured (1:50 dilution) in 50 mL LB with appropriate antibiotics at 37℃ for 1 – 1.5 hour until a maximum OD_600_ of 0.15 was reached. Bacteria were then harvested by centrifugation at 3,000*g* for 5 min, washed with ice-cold PBS three times, and resuspended in ice-cold PBS for injection. A total of 40 μL of each suspension containing 1 × 10^6^ CFU bacteria was injected intratumorally. In the abscopal experiments, bacteria were injected only into the tumors on the right hind flank, leaving the left tumor untreated. For intravenous injection, bacteria were prepared as above described, and 100 uL bacteria suspension (2 × 10^8^ CFU/mL in PBS) was injected via tail vein. Mice were subsequently administered with 500 mg/kg 5-fluorocytosine (5-FC) via intraperitoneal injection according to the indicated schedules.

For the tumor-targeting experiment, mice were sacrificed 5 days post-intravenous injection of EcNx-pCD bacteria (2 × 10^7^ CFU/mouse in 100 uL PBS). Tumor tissues and main organs were collected for CFU determination and WB detection of pCD. Briefly, tumors and organs were weighted and homogenized in 2 mL PBS, and a portion of the homogenized contents were serial-diluted and plated on LB agar plates for CFU analysis. The rest of the homogenized tissues were centrifuged at 12,000 rpm for 15 min, and the supernatants were collected for pCD WB. Supernatants in normalized volume (based on tissue weights) were boiled with SDS loading dye and loaded to a Bolt™ 4-12% Bis-Tris Plus gel for WB as above described.

### Immune phenotyping by flow cytometry

Tumors were treated as illustrated in figure 3A and harvested at indicated time points (day 16 or day 22 post-tumor cell inoculation). Tumor tissues were fragmented and digested in wash media (RPMI-1640 supplemented with 5% FBS, 10 mM HEPES, 1× Glutamax, 1× Pen/Strep) with 1 mg/mL collagenase A and 0.5 μg/mL DNAse I in a 37℃ shaking incubator for 45 minutes. Single cell suspension was prepared by passing the dissociated cell suspension through a 100 μm cell strainer placed atop a 50 mL falcon tube. Cells were then either restimulated for cytokine staining or directly stained for flow cytometry analysis. For *ex vivo* restimulation with PMA/ionomycin, aliquots of single cell suspensions were incubated in 10% complete RPMI-1640 with PMA (50 ng/mL), ionomycin (500 ng/mL) and brefeldin A (1 μg/mL) at 37°C for 3 hours. Live/dead staining was performed using Ghost Dye Red 780 (Tonbo Biosciences), as per the manufacturer’s protocol. Cells were subsequently stained with different antibodies for flow cytometric analyses, with intracellular staining performed using the Foxp3 /Transcription Factor Staining Buffer Kit (Tonbo, Cat. No.: TNB-1020-L050; TNB-1022-L160) as per the manufacturer’s protocol. Antibodies used included anti-CD45 (clone 30-F11, BioLegend), NK1.1 (clone PK136, BD), CD3ε (clone 145-2C11, Tonbo), TCRβ (clone H57-597, BD), CD4 (clone RM4-5, BD), CD8 (clone 53-6.7, Tonbo), Foxp3 (clone FJK-16s, eBioscience), Granzyme B (clone QA16A02 BioLegend), Ki-67 (clone SolA15, eBioscience), TIM-3 (clone RMT3-23, BioLegend), PD-1 (clone 29F.1A12, BioLegend), IFNγ (clone XMG1.2, Tonbo), CD11b (clone M1/70, BD), CD11c (clone HL3, BD), Ly6C (clone HK1.4, BioLegend), Ly6G (clone 1A8, BioLegend), F4/80 (clone BM8.1, Tonbo), CD206 (clone C068C2, BioLegend), MHCII (clone M5/114.15.2, Tonbo), CD103 (clone M290, BD), CD80 (clone 16-10A1, Tonbo), CD86 (clone GL-1, BD), PD-L1 (clone 10F.9G2, Tonbo).

### Statistical analysis

Data are presented as mean ± standard error of the mean. Statistical analysis was performed using GraphPad Prism 10 software. Student’s t-test was performed when only two value sets were compared. One- or two-way ANOVA was performed for the comparisons between multiple groups. A value of p < 0.05 was considered significant. The specific statistical test, statistical significance and the number of samples are noted in figures or figure legends where appropriate.

## Acknowledgments

We thank members of the Arpaia lab for helpful discussions. We thank Florian Hoffmann of the S.H. Sternberg lab for technical support. The flow cytometry experiments were performed in the Flow Cytometry Core Facility of Department of Microbiology and Immunology, Columbia University.

## Author contributions

Z.Y. and N.A. conceived and designed the study with input from T.D.; Z.Y. performed experiments and analyzed data. J.I. made the *preTA* knockout strain. N.C., D.L.M., K.d.l.S.-A., and F.L. assisted with animal studies. Z.Y. and N.A. wrote the manuscript with input from all authors.

## Data and materials availability

All data are available in the main text or the supplementary materials. All inquiries regarding the reagents and resources cited in this work should be directed to N.A., who will address any specific requests for information. Bacterial strains and plasmids are available through N.A., pending scientific review and a completed material transfer agreement with Columbia University. Requests for these strains and plasmids should be submitted to N.A.

## Supplementary Materials

### Supplementary Figures

**Fig. S1.**
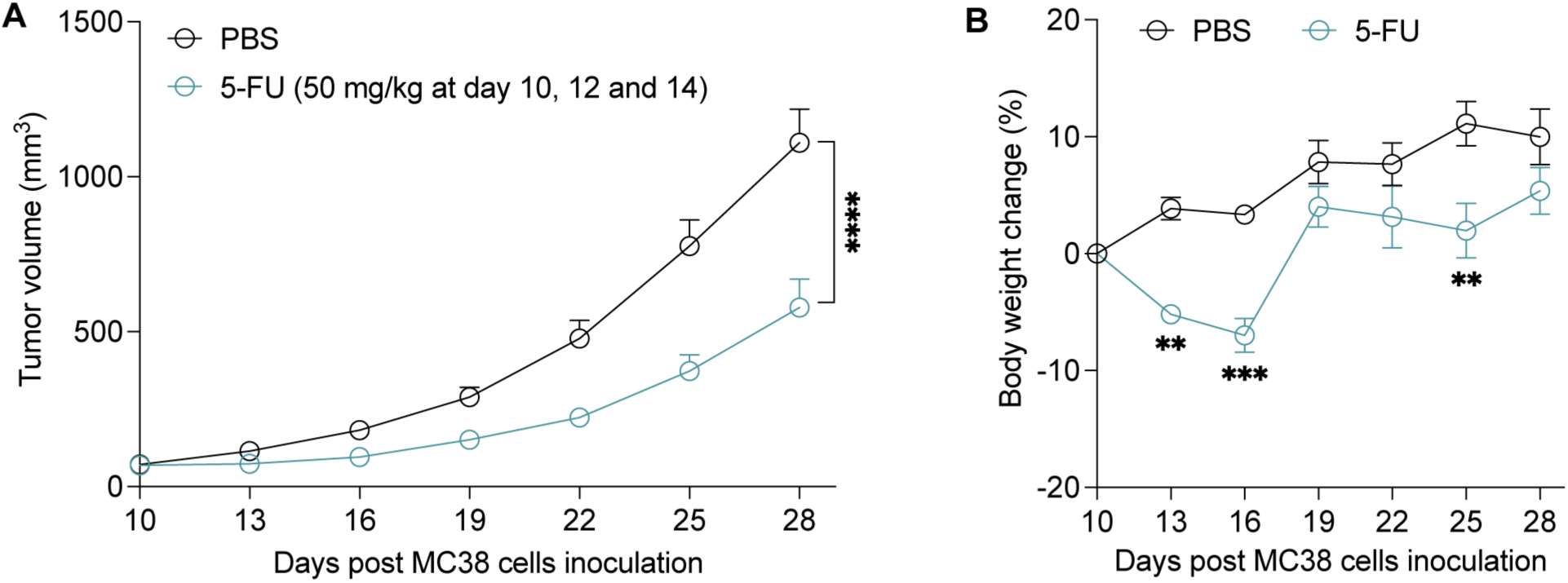
Systemic administration of 5-FU at efficacious dosage causes significant toxicity in mice. **(A)** MC38 tumor growth curves of mice receiving intraperitoneal injections of 50 mg/kg 5-FU at days 10, 12 and 14 post-cancer cell inoculation. **(B)** Body weight change of MC38 tumor-bearing mice during the experimental period.

**Fig. S2.**
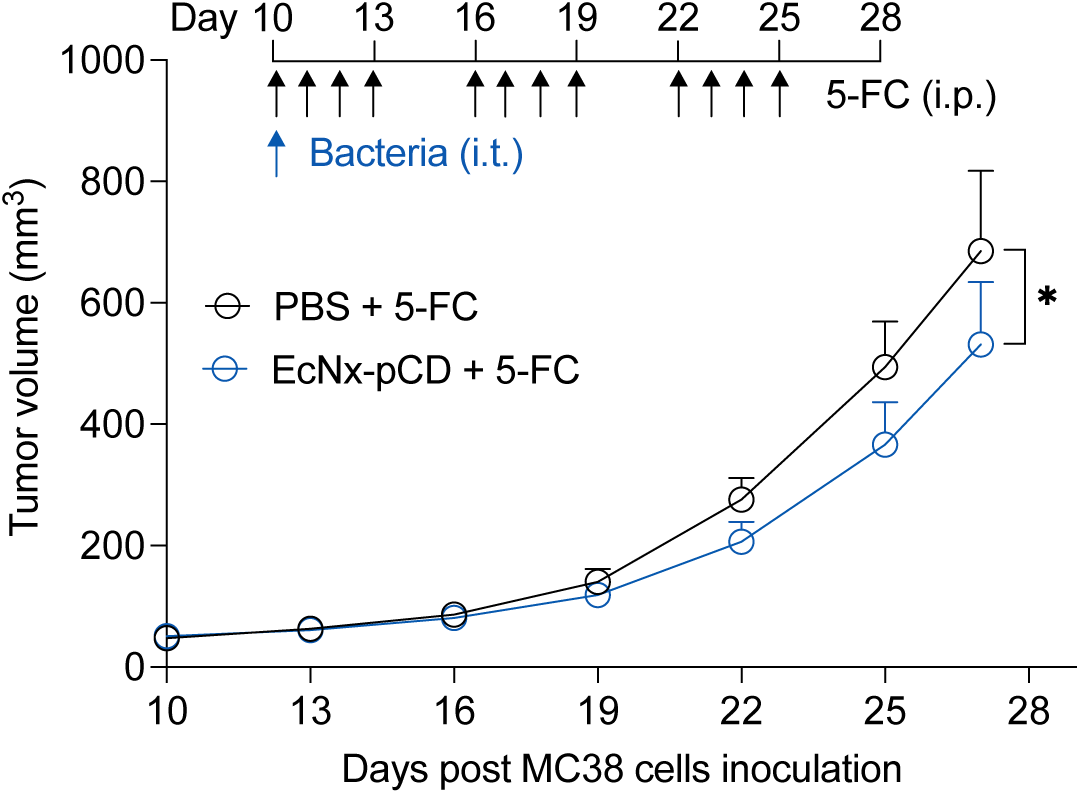
EcNx-pCD bacteria demonstrate minor antitumor effect. MC38 tumor growth curves of mice treated with PBS plus 5-FC (500 mg/kg) or a single injection of EcNx-pCD bacteria (1 × 10^6^ CFU/ tumor) plus 5-FC (500 mg/kg). Data are presented as mean ± sem. * p = 0.0478; Two-way ANOVA with Fisher’s least significant difference (LSD) test.

**Fig. S3.**
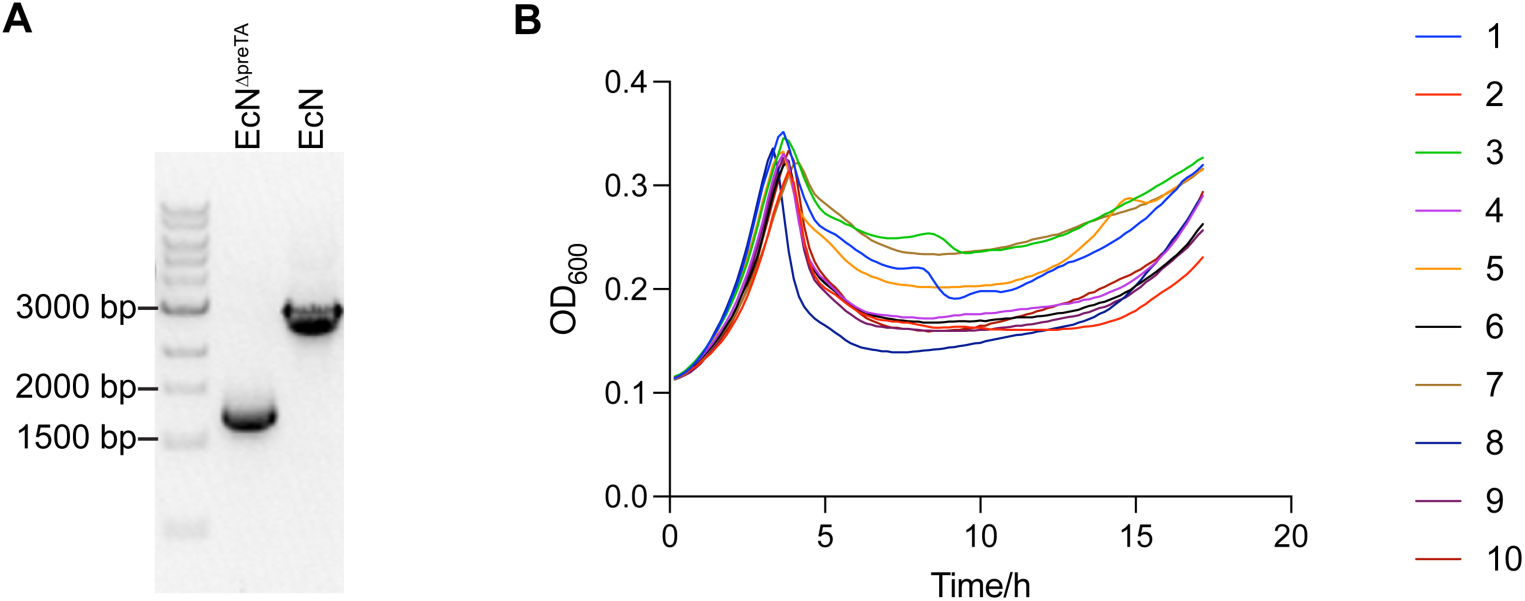
EcNx^ΔpreTA^ strain. **(A)** Colony PCR analysis of EcN^ΔpreTA^ strain to confirm the knockout of *preTA* operon. **(B)** Plate reader experiment showing the bacterial growth dynamics over time of ten EcNx^ΔpreTA^ colonies after the integration of lysing circuit into EcN^ΔpreTA^ bacteria.

**Fig. S4.**
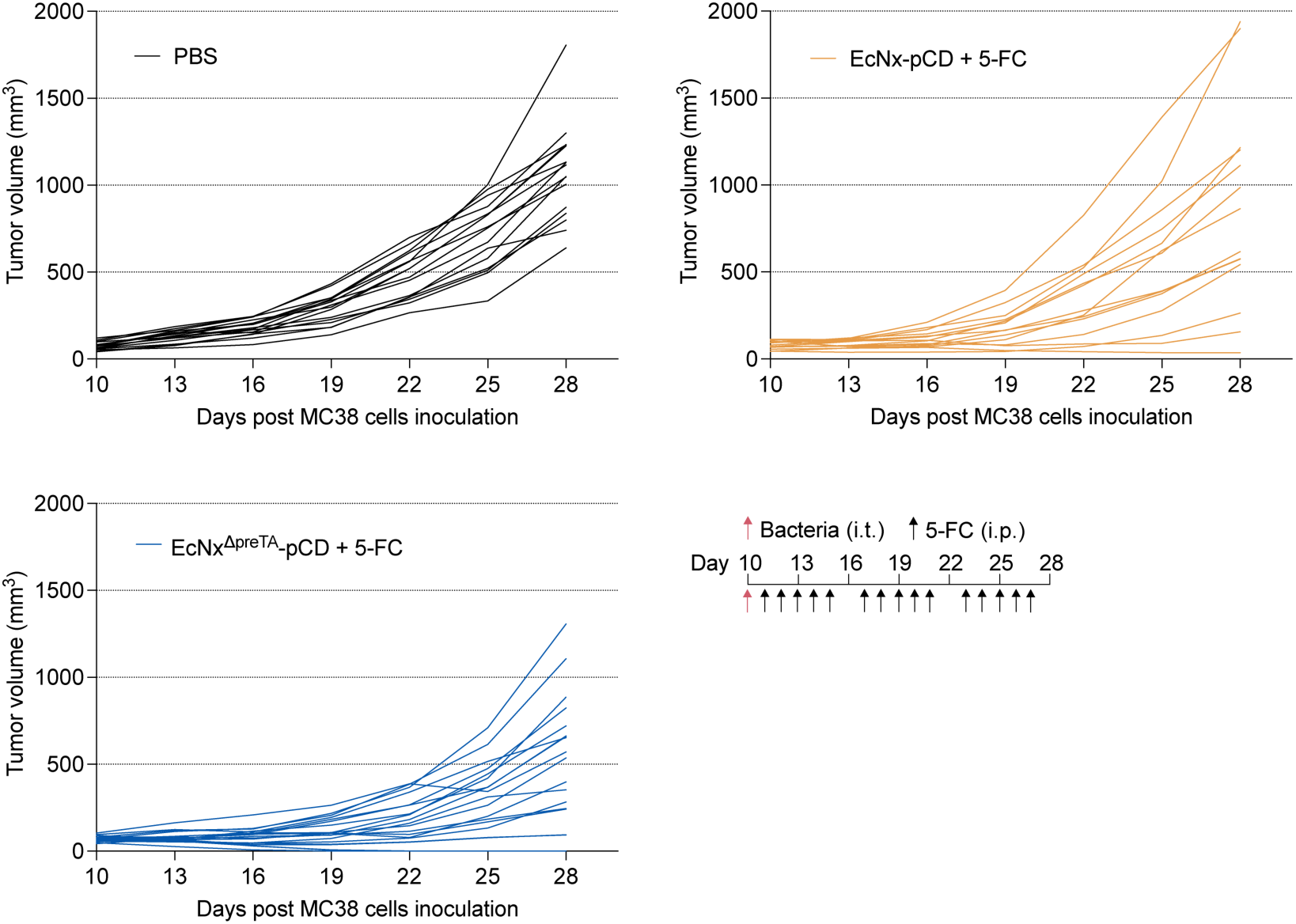
Individual MC38 tumor growth trajectories. Tumor-bearing C57BL/6NJ mice received a single intratumoral injection of the indicated bacteria (1 × 10^6^ CFU/tumor) followed by intraperitoneal injections of 5-FC (500 mg/kg).

**Fig. S5.**
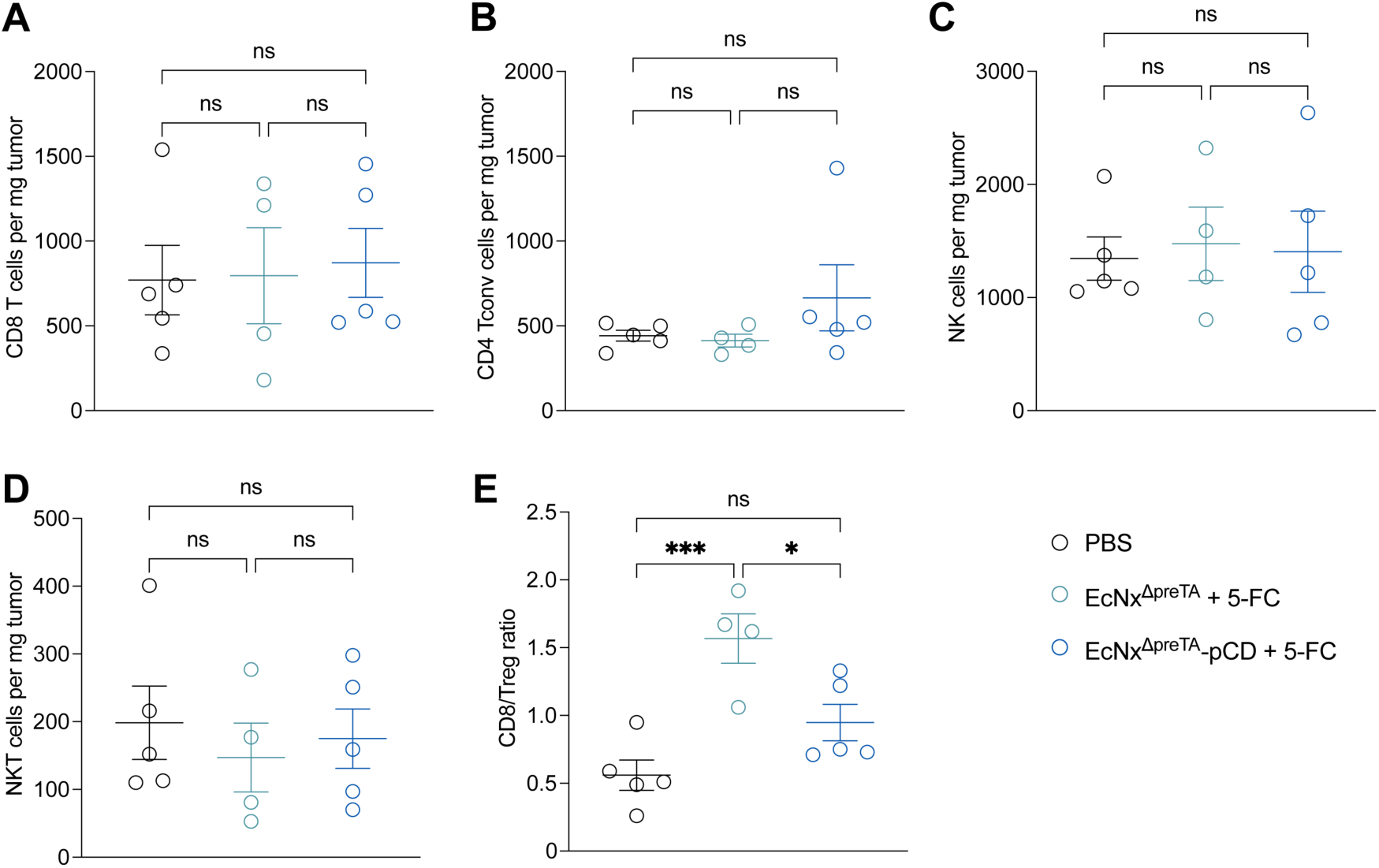
EcNx^ΔpreTA^-pCD/5-FC therapy cannot facilitate tumor infiltration of antitumor effector cells. C57BL/6NJ mice bearing subcutaneous MC38 tumors were treated as indicated in figure 3A. Tumor were collected for flow cytometric analysis of intratumoral immune cells on day 16. Number of **(A)** CD8 T cells (CD45^+^NK1.1^−^CD3^+^TCRβ^+^CD8^+^), **(B)** CD4 Tconv cells (CD45^+^NK1.1^−^CD3^+^TCRβ^+^CD4^+^Foxp3^−^), **(C)** NK cells (CD45^+^NK1.1^+^CD3^−^TCRβ^−^) and **(D)** NKT cells (CD45^+^NK1.1^+^CD3^+^TCRβ^+^) per mg tumor. **(E)** CD8 T cells/Tregs ratio. (A to E) Data are presented as mean ± sem. ns, not significant; * p = 0.0111; *** p = 0.0004; One-way ANOVA with Fisher’s LSD test.

**Fig. S6.**
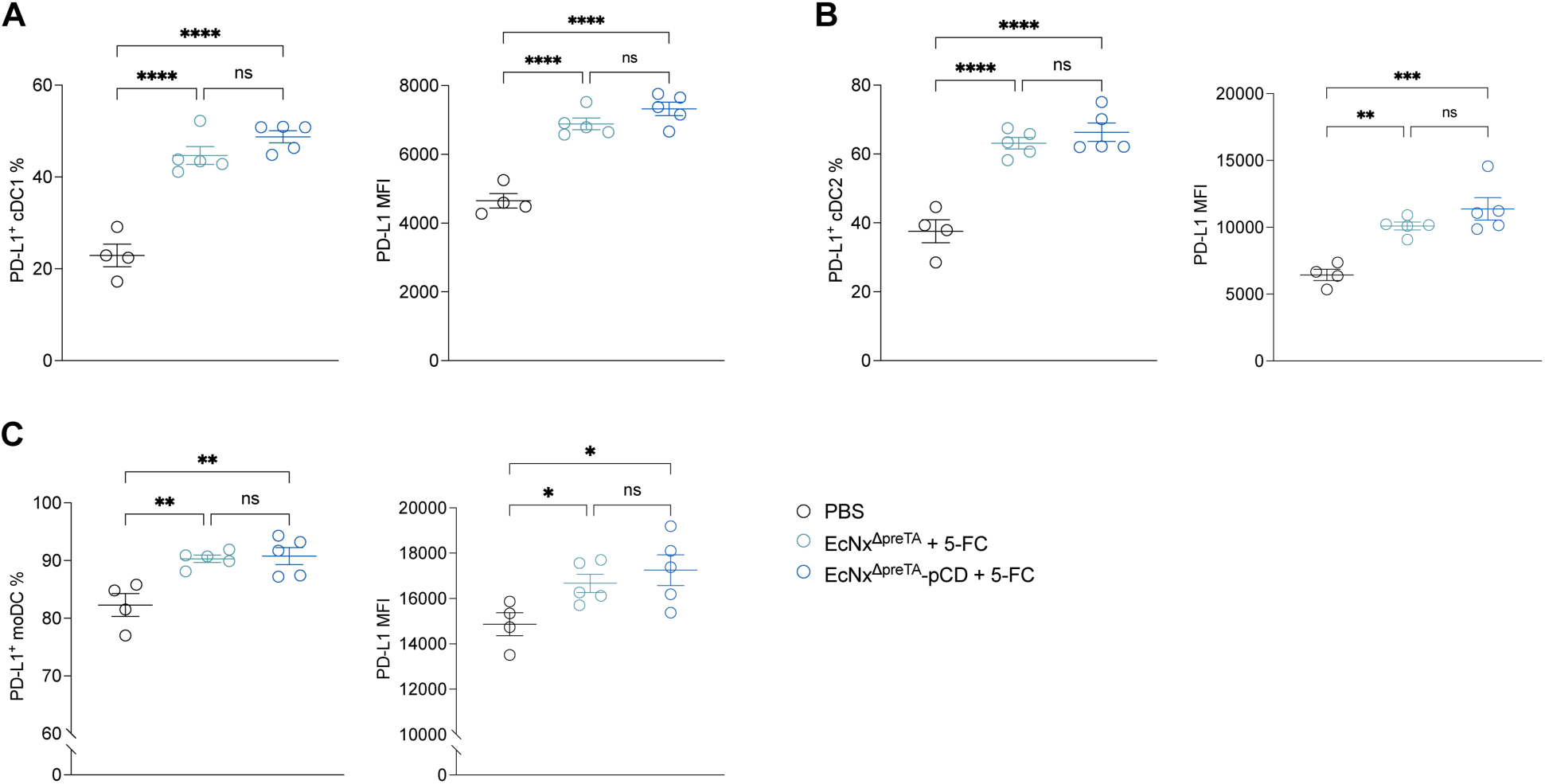
Bacteria treatments upregulate PD-L1 expression on intratumoral dendritic cells. C57BL/6NJ mice bearing subcutaneous MC38 tumors were treated as indicated in figure 3A. Tumor were collected for flow cytometric analysis of intratumoral immune cells on day 22. **(A)** Frequency of PD-L1^+^ cDC1 (CD45^+^Ly6G^−^Ly6C^−^F4/80^−^CD11c^+^MHCII^+^CD103^+^) and PD-L1 MFI of cDC1. **(B)** Frequency of PD-L1^+^ cDC2 (CD45^+^Ly6G^−^Ly6C^−^F4/80^−^CD11c^+^MHCII^+^CD11b^+^CD103^−^) and PD-L1 MFI of cDC2. **(C)** Frequency of PD-L1^+^ moDCs (CD45^+^Ly6G^−^Ly6C^+^CD11c^+^MHCII^+^) and PD-L1 MFI of moDC. (A to C) Data are presented as mean ± sem. ns, not significant; * p < 0.05; ** p < 0.01; *** p < 0.001; **** p < 0.0001; One-way ANOVA with Fisher’s LSD test.

**Fig. S7.**
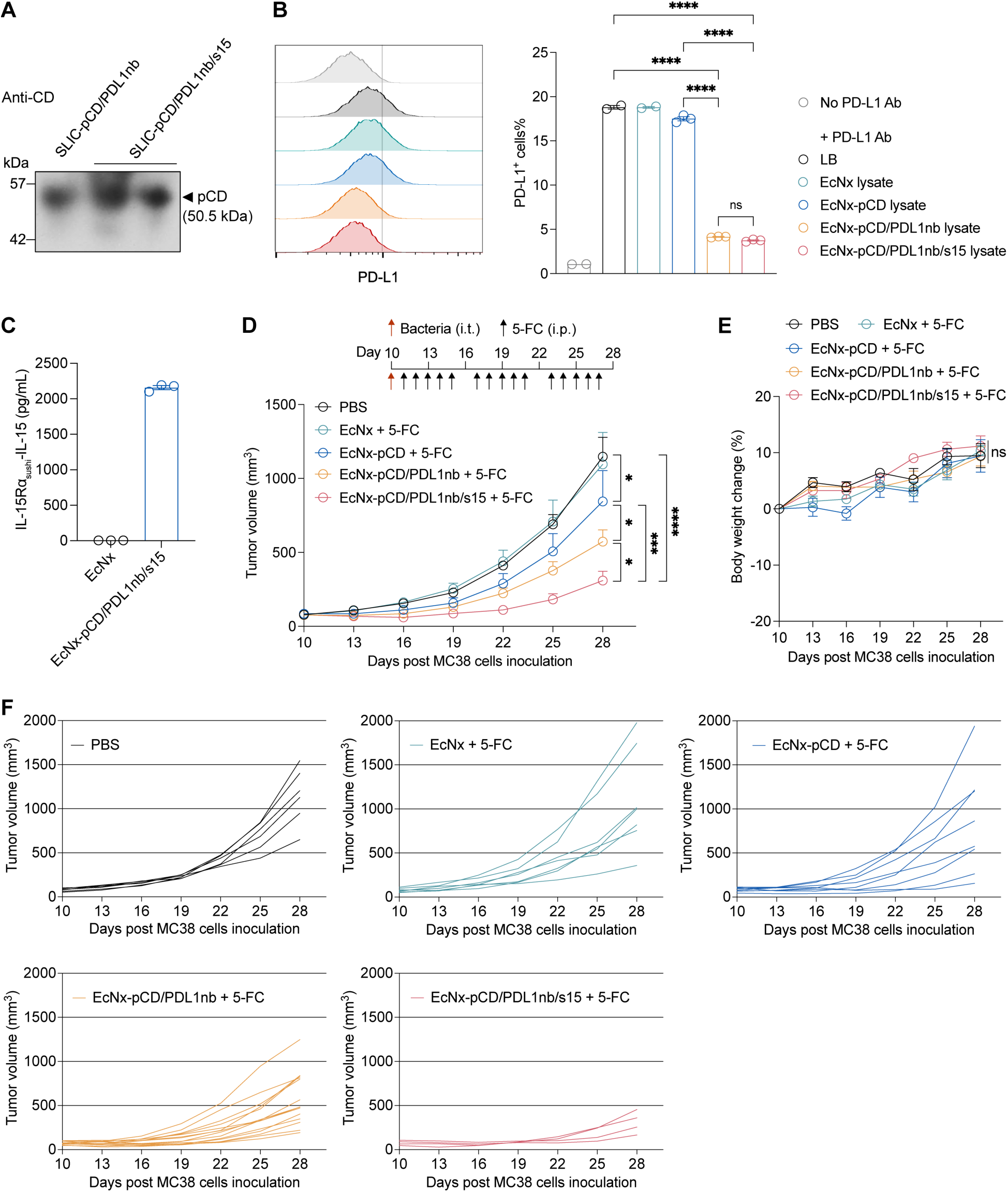
EcNx bacteria co-expressing pCD, PDL1nb and s15 enable efficient chemoimmunotherapy. **(A)** Western blot image showing the expression of pCD by EcNx-pCD/PDL1nb and EcNx-pCD/PDL1nb/s15 strains. **(B)** Merged flow cytometric histogram and percentage of PD-L1^+^ MC38 cells showing the bacterial production and activity of PDL1nb. MC38 cells were pre-incubated with lysates of indicated strains for 20 min and then stained with APC-labeled PD-L1 antibody for flow cytometric analysis. Data are presented as mean ± sem; ns, not significant; **** p<0.0001; One-way ANOVA with Tukey’s multiple comparisons test. **(C)** ELISA quantification of IL-15Rα_sushi_-IL-15 concentration in the lysates of EcNx and EcNx-pCD/PDL1nb/s15 bacteria. Data are presented as mean ± sem. **(D)** MC38 tumor growth curves from C57BL/6NJ mice that received intratumoral injection of the indicated bacteria (red arrow; 1 × 10^6^ CFU/tumor) and intraperitoneal injections of 5-FC (black arrows; 500 mg/kg). **(E)** Body weight change of MC38 tumor-bearing mice during the treatment period. **(F)** Individual MC38 tumor growth trajectories. (D to F) Data are combined from 2 independent experiments and presented as mean ± sem (n ≥ 4 tumors per group); (E) * p<0.05; *** p< 0.001; ****P < 0.0001; Two-way ANOVA with Holm-Šídák multiple comparisons test.

**Fig. S8.**
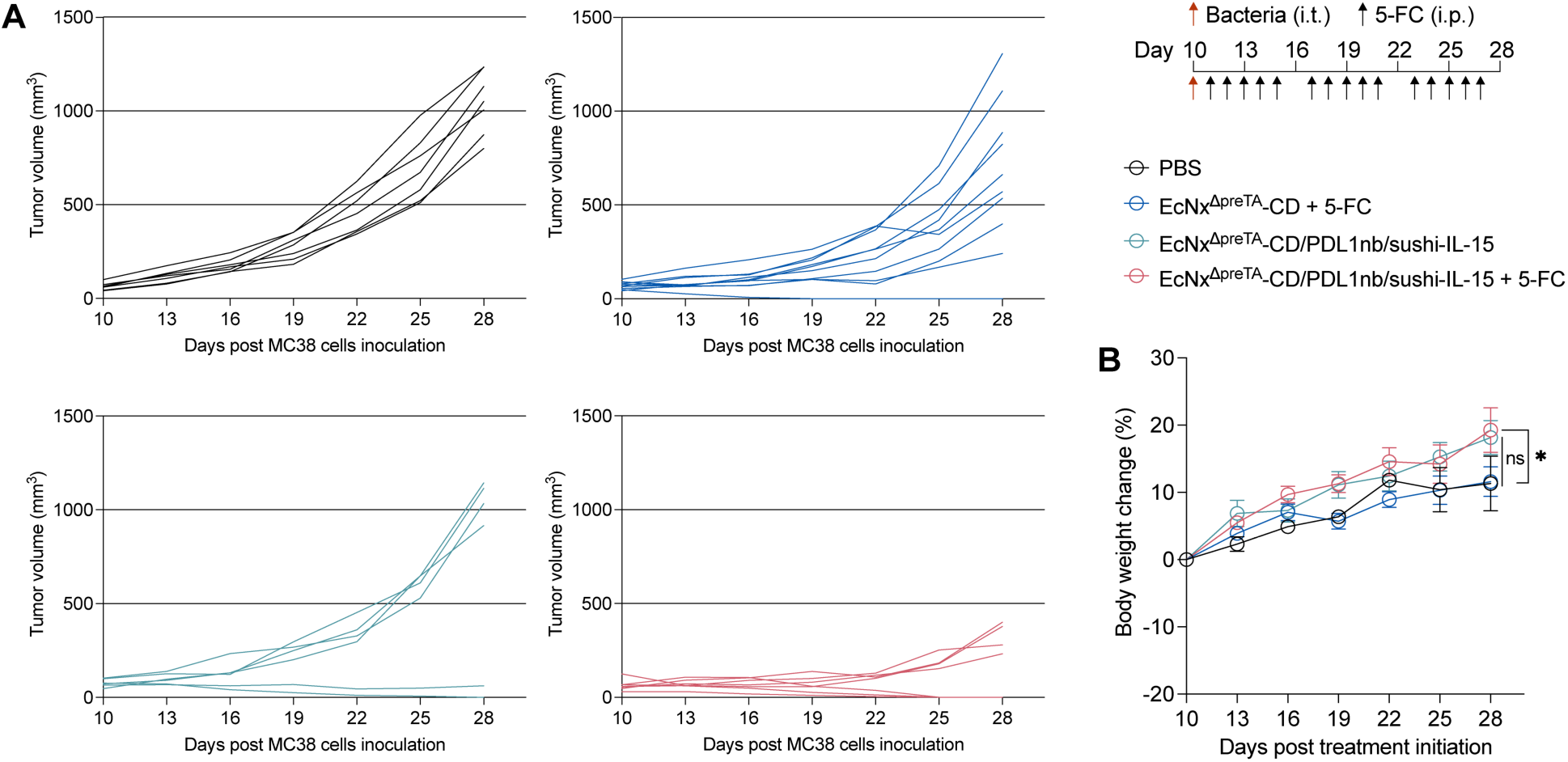
**(A)** Individual MC38 tumor growth trajectories and **(B)** body weight changes of mice that received intratumoral injection of the indicated bacteria (1 × 10^6^ CFU/tumor) on day 10 post-subcutaneous inoculation of MC38 cells and following intraperitoneal injections of 5-FC (500 mg/kg) as illustrated. (B) Data are presented as mean ± sem. ns, not significant; * p < 0.05; Two-way ANOVA with Holm-Šídák multiple comparisons test.

**Fig. S9.**
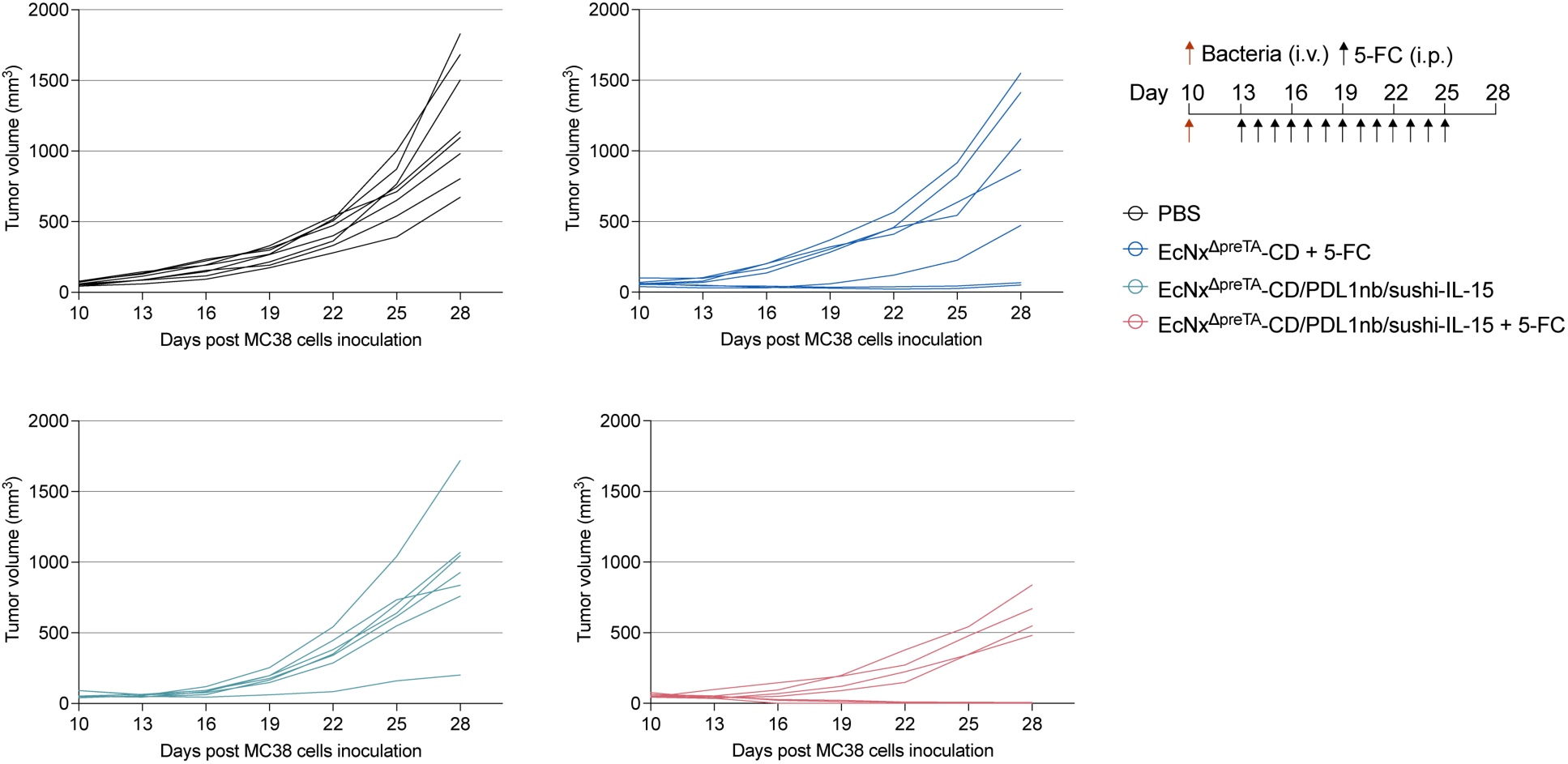
Individual MC38 tumor growth trajectories. MC38 tumor-bearing mice received intravenous injection of indicated bacteria (2 × 10^7^ CFU/mouse) on day 10 post-subcutaneous inoculation of MC38 cells and intraperitoneal injections of 5-FC (500 mg/kg) as illustrated.

**Fig. S10.**
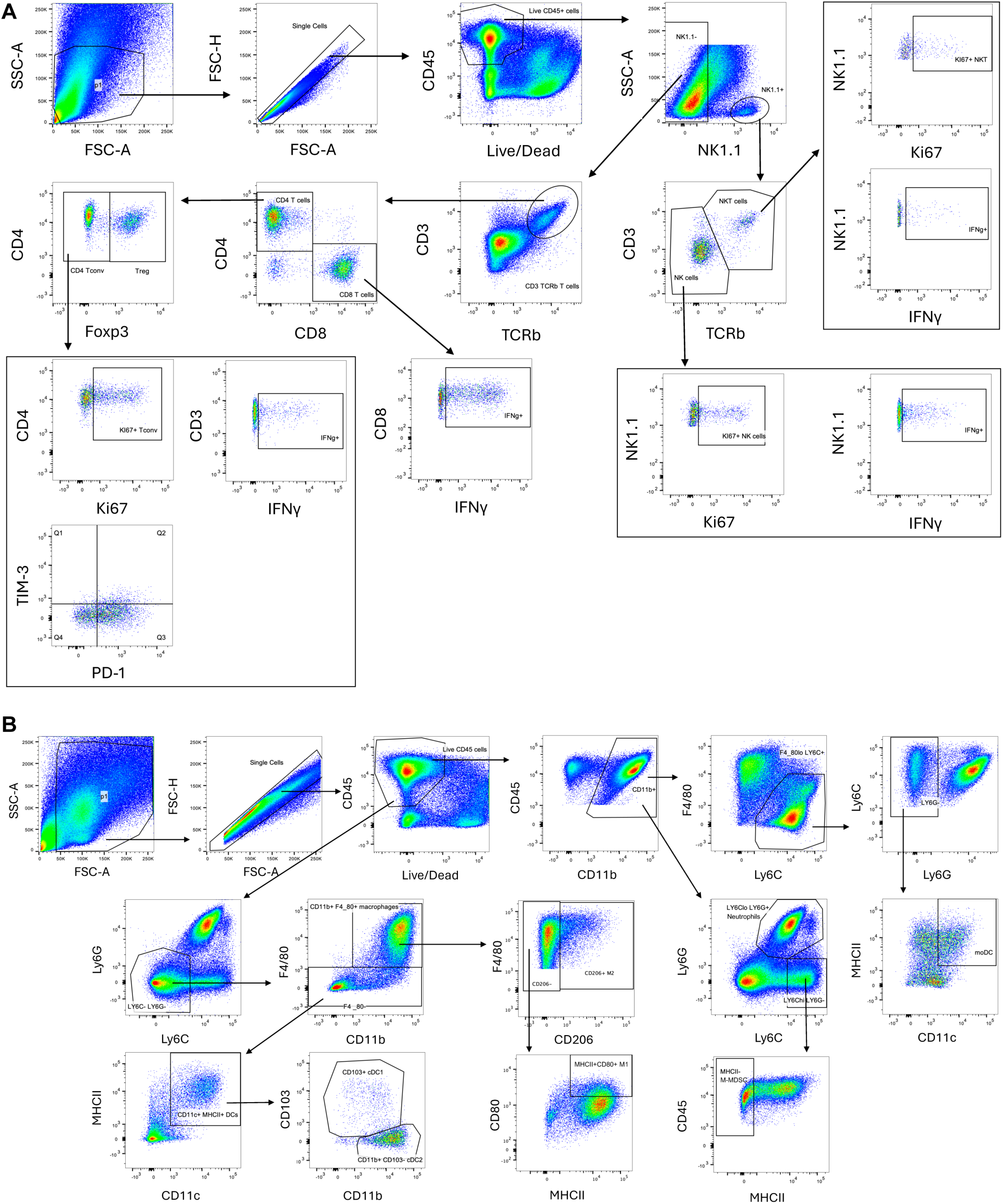
Gating strategies for flow cytometric analyses of intratumoral **(A)** lymphocytes and **(B)** myeloid cells.

**Fig. S11.**
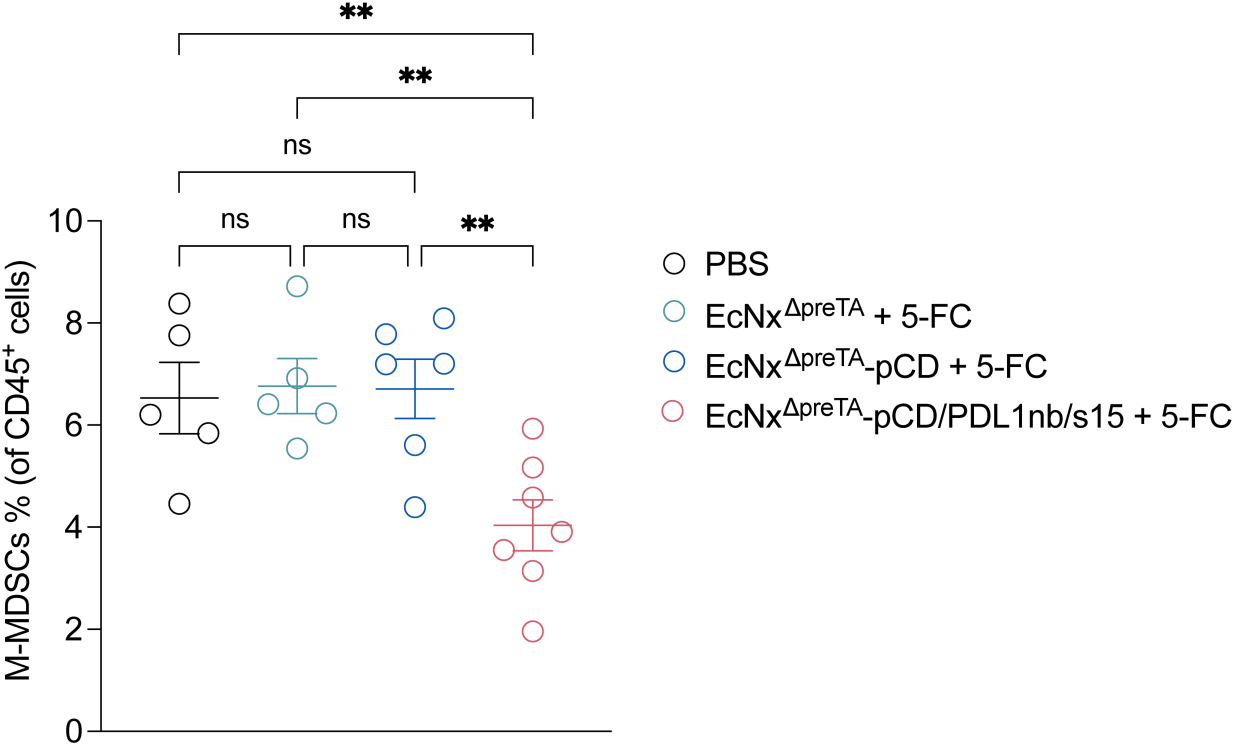
Frequency of M-MDSCs of CD45^+^ cells. Mice bearing subcutaneous MC38 tumors received intratumoral injection of either EcNx^ΔpreTA^, EcNx^ΔpreTA^-pCD, or EcNx^ΔpreTA^-pCD/PDL1nb/s15 bacteria (1 × 10^6^ CFU/tumor) and intraperitoneal injections of 5-FC (500 mg/kg) as illustrated in figure 5A. Tumors were collected on day 22 post cancer cell inoculation for flow cytometric analysis of intratumoral immune cells. M-MDSCs, Monocytic Myeloid-Derived Suppressor Cells (CD45^+^CD11b^+^Ly6G^−^Ly6C^hi^MHCII^−^). Data are presented as mean ± sem. ns, not significant; ** p < 0.01; One-way ANOVA with Fisher’s LSD test.

**Fig. S12.**
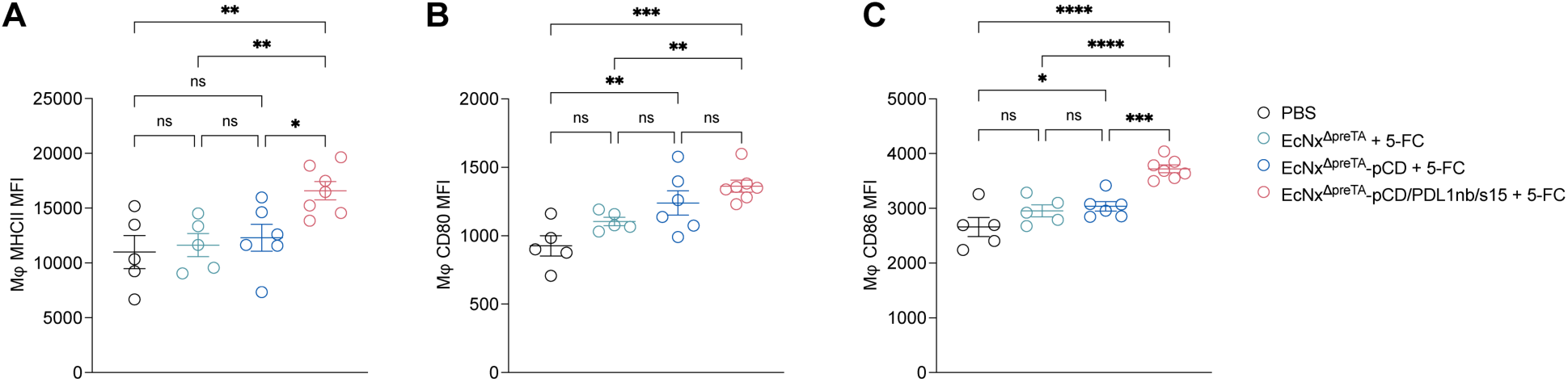
MFIs showing the expression of MHCII, CD80 and CD86 on intratumoral macrophages. **(A)** MHCII. **(B)** CD80. **(C)** CD86. Mice bearing subcutaneous MC38 tumors received intratumoral injection of either EcNx^ΔpreTA^, EcNx^ΔpreTA^-pCD, or EcNx^ΔpreTA^-pCD/PDL1nb/s15 bacteria (1 × 10^6^ CFU/tumor) and intraperitoneal injections of 5-FC (500 mg/kg) as illustrated in figure 5A. Tumors were collected on day 22 post-cancer cell inoculation for flow cytometric analysis of intratumoral immune cells. Mφ, macrophages, CD45^+^Ly6G^−^Ly6C^−^CD11b^+^F4/80^+^. Data are presented as mean ± sem. ns, not significant; * p < 0.05; ** p < 0.01; *** p < 0.001; **** p < 0.0001; One-way ANOVA with Fisher’s LSD test.

## References

1. U. Anand, A. Dey, A. K. S. Chandel, R. Sanyal, A. Mishra, D. K. Pandey, V. De Falco, A. Upadhyay, R. Kandimalla, A. Chaudhary, J. K. Dhanjal, S. Dewanjee, J. Vallamkondu, J. M. Pérez de la Lastra, Cancer chemotherapy and beyond: Current status, drug candidates, associated risks and progress in targeted therapeutics. Genes Dis. 10, 1367–1401 (2023).

2. R. Mooney, A. Abdul Majid, J. Batalla, A. J. Annala, K. S. Aboody, Cell-mediated enzyme prodrug cancer therapies. Adv. Drug Delivery Rev. 118, 35–51 (2017).

3. G. Xu, H. L. McLeod, Strategies for Enzyme/Prodrug Cancer Therapy. Clin. Cancer Res. 7, 3314–3324 (2001).

4. D. B. Longley, D. P. Harkin, P. G. Johnston, 5-Fluorouracil: mechanisms of action and clinical strategies. Nat. Rev. Cancer 3, 330–338 (2003).

5. I. Dagogo-Jack, A. T. Shaw, Tumour heterogeneity and resistance to cancer therapies. Nat. Rev. Clin. Oncol. 15, 81–94 (2018).

6. D. Longley, P. Johnston, Molecular mechanisms of drug resistance. J. Pathol. 205, 275–292 (2005).

7. S. Farkona, E. P. Diamandis, I. M. Blasutig, Cancer immunotherapy: the beginning of the end of cancer? BMC Med. 14, 73 (2016).

8. S. Das, D. B. Johnson, Immune-related adverse events and anti-tumor efficacy of immune checkpoint inhibitors. J. Immunother. Cancer 7, 306 (2019).

9. Z. Yang, J. K.-L. Sun, M. M. Lee, M. K. Chan, Restoration of p53 activity via intracellular protein delivery sensitizes triple negative breast cancer to anti-PD-1 immunotherapy. J. Immunother. Cancer 10, e005068 (2022).

10. L. Galluzzi, A. Buqué, O. Kepp, L. Zitvogel, G. Kroemer, Immunological Effects of Conventional Chemotherapy and Targeted Anticancer Agents. Cancer Cell 28, 690–714 (2015).

11. L. Zitvogel, L. Apetoh, F. Ghiringhelli, G. Kroemer, Immunological aspects of cancer chemotherapy. Nat. Rev. Immunol. 8, 59–73 (2008).

12. L. Galluzzi, J. Humeau, A. Buqué, L. Zitvogel, G. Kroemer, Immunostimulation with chemotherapy in the era of immune checkpoint inhibitors. Nat. Rev. Clin. Oncol. 17, 725–741 (2020).

13. J. Tian, D. Zhang, V. Kurbatov, Q. Wang, Y. Wang, D. Fang, L. Wu, M. Bosenberg, M. D. Muzumdar, S. Khan, Q. Lu, Q. Yan, J. Lu, 5-Fluorouracil efficacy requires anti-tumor immunity triggered by cancer-cell-intrinsic STING. EMBO J. 40, e106065 (2021).

14. M. Yi, X. Zheng, M. Niu, S. Zhu, H. Ge, K. Wu, Combination strategies with PD-1/PD-L1 blockade: current advances and future directions. Mol. Cancer 21, 28 (2022).

15. K. M. Heinhuis, W. Ros, M. Kok, N. Steeghs, J. H. Beijnen, J. H. M. Schellens, Enhancing antitumor response by combining immune checkpoint inhibitors with chemotherapy in solid tumors. Ann. Oncol. 30, 219–235 (2019).

16. D. Salas-Benito, J. L. Pérez-Gracia, M. Ponz-Sarvisé, M. E. Rodriguez-Ruiz, I. Martínez-Forero, E. Castañón, J. M. López-Picazo, M. F. Sanmamed, I. Melero, Paradigms on Immunotherapy Combinations with Chemotherapy. Cancer Discovery 11, 1353–1367 (2021).

17. P. Gotwals, S. Cameron, D. Cipolletta, V. Cremasco, A. Crystal, B. Hewes, B. Mueller, S. Quaratino, C. Sabatos-Peyton, L. Petruzzelli, J. A. Engelman, G. Dranoff, Prospects for combining targeted and conventional cancer therapy with immunotherapy. Nat. Rev. Cancer 17, 286–301 (2017).

18. J.-J. Min, H.-J. Kim, J. H. Park, S. Moon, J. H. Jeong, Y.-J. Hong, K.-O. Cho, J. H. Nam, N. Kim, Y.-K. Park, H.-S. Bom, J. H. Rhee, H. E. Choy, Noninvasive Real-time Imaging of Tumors and Metastases Using Tumor-targeting Light-emitting Escherichia coli. Mol. Imaging Biol. 10, 54–61 (2008).

19. R. L. Vincent, C. R. Gurbatri, F. Li, A. Vardoshvili, C. Coker, J. Im, E. R. Ballister, M. Rouanne, T. Savage, K. de los Santos-Alexis, A. Redenti, L. Brockmann, M. Komaranchath, N. Arpaia, T. Danino, Probiotic-guided CAR-T cells for solid tumor targeting. Science 382, 211–218 (2023).

20. C. R. Gurbatri, I. Lia, R. Vincent, C. Coker, S. Castro, P. M. Treuting, T. E. Hinchliffe, N. Arpaia, T. Danino, Engineered probiotics for local tumor delivery of checkpoint blockade nanobodies. Sci. Transl. Med. 12, eaax0876 (2020).

21. T. M. Savage, R. L. Vincent, S. S. Rae, L. H. Huang, A. Ahn, K. Pu, F. Li, K. de los Santos-Alexis, C. Coker, T. Danino, N. Arpaia, Chemokines expressed by engineered bacteria recruit and orchestrate antitumor immunity. Sci. Adv. 9, eadc9436 (2023).

22. S. Chowdhury, S. Castro, C. Coker, T. E. Hinchliffe, N. Arpaia, T. Danino, Programmable bacteria induce durable tumor regression and systemic antitumor immunity. Nat. Med. 25, 1057–1063 (2019).

23. F. Li, Z. Yang, T. M. Savage, R. L. Vincent, K. de los Santos-Alexis, A. Ahn, M. Rouanne, D. L. Mariuzza, T. Danino, N. Arpaia, Programmable bacteria synergize with PD-1 blockade to overcome cancer cell–intrinsic immune resistance mechanisms. Sci. Immunol. 9, eadn9879 (2024).

24. A. B. P. van Kuilenburg, Dihydropyrimidine dehydrogenase and the efficacy and toxicity of 5-fluorouracil. Eur. J. Cancer 40, 939–950 (2004).

25. B. Gustavsson, G. Carlsson, D. Machover, N. Petrelli, A. Roth, H.-J. Schmoll, K.-M. Tveit, F. Gibson, A Review of the Evolution of Systemic Chemotherapy in the Management of Colorectal Cancer. Clin. Colorectal Cancer 14, 1–10 (2015).

26. M. O. Din, T. Danino, A. Prindle, M. Skalak, J. Selimkhanov, K. Allen, E. Julio, E. Atolia, L. S. Tsimring, S. N. Bhatia, J. Hasty, Synchronized cycles of bacterial lysis for in vivo delivery. Nature 536, 81–85 (2016).

27. M. M. Martino, P. S. Briquez, E. Güç, F. Tortelli, W. W. Kilarski, S. Metzger, J. J. Rice, G. A. Kuhn, R. Müller, M. A. Swartz, J. A. Hubbell, Growth Factors Engineered for Super-Affinity to the Extracellular Matrix Enhance Tissue Healing. Science 343, 885–888 (2014).

28. M. Fuchita, A. Ardiani, L. Zhao, K. Serve, B. L. Stoddard, M. E. Black, Bacterial Cytosine Deaminase Mutants Created by Molecular Engineering Show Improved 5-Fluorocytosine–Mediated Cell Killing In vitro and In vivo. Cancer Res. 69, 4791–4799 (2009).

29. R. Hidese, H. Mihara, T. Kurihara, N. Esaki, Escherichia coli Dihydropyrimidine Dehydrogenase Is a Novel NAD-Dependent Heterotetramer Essential for the Production of 5,6-Dihydrouracil. J. Bacteriol. 193, 989–993 (2011).

30. T. T. M. Nguyen, V.-H. Mai, H. S. Kim, D. Kim, M. Seo, Y. J. An, S. Park, Real-Time Monitoring of Host–Gut Microbial Interspecies Interaction in Anticancer Drug Metabolism. J. Am. Chem. Soc. 144, 8529–8535 (2022).

31. K. D. LaCourse, M. Zepeda-Rivera, A. G. Kempchinsky, A. Baryiames, S. S. Minot, C. D. Johnston, S. Bullman, The cancer chemotherapeutic 5-fluorouracil is a potent Fusobacterium nucleatum inhibitor and its activity is modified by intratumoral microbiota. Cell Rep. 41, 111625 (2022).

32. P. Spanogiannopoulos, T. S. Kyaw, B. G. H. Guthrie, P. H. Bradley, J. V. Lee, J. Melamed, Y. N. A. Malig, K. N. Lam, D. Gempis, M. Sandy, W. Kidder, E. L. Van Blarigan, C. E. Atreya, A. Venook, R. R. Gerona, A. Goga, K. S. Pollard, P. J. Turnbaugh, Host and gut bacteria share metabolic pathways for anti-cancer drug metabolism. Nat. Microbiol. 7, 1605–1620 (2022).

33. A. Galetto, S. Buttiglieri, S. Forno, F. Moro, A. Mussa, L. Matera, Drug- and cell-mediated antitumor cytotoxicities modulate cross-presentation of tumor antigens by myeloid dendritic cells. Anti-Cancer Drugs 14, 833 (2003).

34. J. Vincent, G. Mignot, F. Chalmin, S. Ladoire, M. Bruchard, A. Chevriaux, F. Martin, L. Apetoh, C. Rébé, F. Ghiringhelli, 5-Fluorouracil selectively kills tumor-associated myeloid-derived suppressor cells resulting in enhanced T cell-dependent antitumor immunity. Cancer Res. 70, 3052–3061 (2010).

35. L. Van Der Kraak, G. Goel, K. Ramanan, C. Kaltenmeier, L. Zhang, D. P. Normolle, G. J. Freeman, D. Tang, K. S. Nason, J. M. Davison, J. D. Luketich, R. Dhupar, M. T. Lotze, 5-Fluorouracil upregulates cell surface B7-H1 (PD-L1) expression in gastrointestinal cancers. J. Immunother. Cancer 4, 65 (2016).

36. Y. Wu, Z. Deng, H. Wang, W. Ma, C. Zhou, S. Zhang, Repeated cycles of 5-fluorouracil chemotherapy impaired anti-tumor functions of cytotoxic T cells in a CT26 tumor-bearing mouse model. BMC Immunol. 17, 29 (2016).

37. X. Hong, T. Dong, T. Yi, J. Hu, Z. Zhang, S. Lin, W. Niu, Department of General Surgery, Zhongshan Hospital, Fudan University, Shanghai 200032, China, Impact of 5-Fu/oxaliplatin on mouse dendritic cells and synergetic effect with a colon cancer vaccine. Chin. J. Cancer Res. 30, 197–208 (2018).

38. S. Orecchioni, G. Talarico, V. Labanca, A. Calleri, P. Mancuso, F. Bertolini, Vinorelbine, cyclophosphamide and 5-FU effects on the circulating and intratumoural landscape of immune cells improve anti-PD-L1 efficacy in preclinical models of breast cancer and lymphoma. Br. J. Cancer 118, 1329–1336 (2018).

39. A. Redenti, J. Im, B. Redenti, F. Li, M. Rouanne, Z. Sheng, W. Sun, C. R. Gurbatri, S. Huang, M. Komaranchath, Y. Jang, J. Hahn, E. R. Ballister, R. L. Vincent, A. Vardoshivilli, T. Danino, N. Arpaia, Probiotic neoantigen delivery vectors for precision cancer immunotherapy. Nature, 1–9 (2024).

40. B. Sun, M. Liu, M. Cui, T. Li, Granzyme B-expressing treg cells are enriched in colorectal cancer and present the potential to eliminate autologous T conventional cells. Immunol. Lett. 217, 7–14 (2020).

41. X. Cao, S. F. Cai, T. A. Fehniger, J. Song, L. I. Collins, D. R. Piwnica-Worms, T. J. Ley, Granzyme B and Perforin Are Important for Regulatory T Cell-Mediated Suppression of Tumor Clearance. Immunity 27, 635–646 (2007).

42. A. J. Kassianos, M. Y. Hardy, X. Ju, D. Vijayan, Y. Ding, A. J. E. Vulink, K. J. McDonald, S. L. Jongbloed, R. B. Wadley, C. Wells, D. N. J. Hart, K. J. Radford, Human CD1c (BDCA-1)+ myeloid dendritic cells secrete IL-10 and display an immuno-regulatory phenotype and function in response to Escherichia coli. Eur. J. Immunol. 42, 1512–1522 (2012).

43. D. J. Propper, F. R. Balkwill, Harnessing cytokines and chemokines for cancer therapy. Nat. Rev. Clin. Oncol. 19, 237–253 (2022).

44. S. K. Perna, B. De Angelis, D. Pagliara, S. T. Hasan, L. Zhang, A. Mahendravada, H. E. Heslop, M. K. Brenner, C. M. Rooney, G. Dotti, B. Savoldo, Interleukin 15 Provides Relief to CTLs from Regulatory T Cell–Mediated Inhibition: Implications for Adoptive T Cell–Based Therapies for Lymphoma. Clin. Cancer Res. 19, 106–117 (2013).

45. M. B. Ahmed, N. Belhadj Hmida, N. Moes, S. Buyse, M. Abdeladhim, H. Louzir, N. Cerf-Bensussan, IL-15 Renders Conventional Lymphocytes Resistant to Suppressive Functions of Regulatory T Cells through Activation of the Phosphatidylinositol 3-Kinase Pathway. J. Immunol. 182, 6763–6770 (2009).

46. E. Mortier, A. Quéméner, P. Vusio, I. Lorenzen, Y. Boublik, J. Grötzinger, A. Plet, Y. Jacques, Soluble Interleukin-15 Receptor α (IL-15Rα)-sushi as a Selective and Potent Agonist of IL-15 Action through IL-15Rβ/γ: HYPERAGONIST IL-15·IL-15Rα FUSION PROTEINS*. J. Biol. Chem. 281, 1612–1619 (2006).

47. F. Veglia, E. Sanseviero, D. I. Gabrilovich, Myeloid-derived suppressor cells in the era of increasing myeloid cell diversity. Nat. Rev. Immunol. 21, 485–498 (2021).

48. S. Jhunjhunwala, C. Hammer, L. Delamarre, Antigen presentation in cancer: insights into tumour immunogenicity and immune evasion. Nat. Rev. Cancer 21, 298–312 (2021).

49. E. Montauti, D. Y. Oh, L. Fong, CD4+ T cells in antitumor immunity. Trends Cancer 0 (2024), doi:10.1016/j.trecan.2024.07.009.

50. J. Borst, T. Ahrends, N. Bąbała, C. J. M. Melief, W. Kastenmüller, CD4+ T cell help in cancer immunology and immunotherapy. Nat. Rev. Immunol. 18, 635–647 (2018).

51. S. T. Ferris, V. Durai, R. Wu, D. J. Theisen, J. P. Ward, M. D. Bern, J. T. Davidson, P. Bagadia, T. Liu, C. G. Briseño, L. Li, W. E. Gillanders, G. F. Wu, W. M. Yokoyama, T. L. Murphy, R. D. Schreiber, K. M. Murphy, cDC1 prime and are licensed by CD4+ T cells to induce anti-tumour immunity. Nature 584, 624–629 (2020).

52. A. M. Miggelbrink, J. D. Jackson, S. J. Lorrey, E. S. Srinivasan, J. Waibl-Polania, D. S. Wilkinson, P. E. Fecci, CD4 T-Cell Exhaustion: Does It Exist and What Are Its Roles in Cancer? Clin. Cancer Res. 27, 5742–5752 (2021).

53. S. Sheikh, D. Ernst, A. Keating, Prodrugs and prodrug-activated systems in gene therapy. Mol. Ther. 29, 1716–1728 (2021).

54. C. R. Gurbatri, G. A. Radford, L. Vrbanac, J. Im, E. M. Thomas, C. Coker, S. R. Taylor, Y. Jang, A. Sivan, K. Rhee, A. A. Saleh, T. Chien, F. Zandkarimi, I. Lia, T. R. M. Lannagan, T. Wang, J. A. Wright, H. Kobayashi, J. Q. Ng, M. Lawrence, T. Sammour, M. Thomas, M. Lewis, L. Papanicolas, J. Perry, T. Fitzsimmons, P. Kaazan, A. Lim, A. M. Stavropoulos, D. A. Gouskos, J. Marker, C. Ostroff, G. Rogers, N. Arpaia, D. L. Worthley, S. L. Woods, T. Danino, Engineering tumor-colonizing E. coli Nissle 1917 for detection and treatment of colorectal neoplasia. Nat. Commun. 15, 646 (2024).

55. J. J. Luke, S. A. Piha-Paul, T. Medina, C. F. Verschraegen, M. Varterasian, A. M. Brennan, R. J. Riese, A. Sokolovska, J. Strauss, D. L. Hava, F. Janku, Phase I Study of SYNB1891, an Engineered E. coli Nissle Strain Expressing STING Agonist, with and without Atezolizumab in Advanced Malignancies. Clin. Cancer Res., OF1–OF10 (2023).

56. J. Nemunaitis, C. Cunningham, N. Senzer, J. Kuhn, J. Cramm, C. Litz, R. Cavagnolo, A. Cahill, C. Clairmont, M. Sznol, Pilot trial of genetically modified, attenuated Salmonella expressing the E. coli cytosine deaminase gene in refractory cancer patients. Cancer Gene Ther. 10, 737–744 (2003).

57. D. Mathios, J. E. Kim, A. Mangraviti, J. Phallen, C.-K. Park, C. M. Jackson, T. Garzon-Muvdi, E. Kim, D. Theodros, M. Polanczyk, A. M. Martin, I. Suk, X. Ye, B. Tyler, C. Bettegowda, H. Brem, D. M. Pardoll, M. Lim, Anti–PD-1 antitumor immunity is enhanced by local and abrogated by systemic chemotherapy in GBM. Sci. Transl. Med. 8, 370ra180–370ra180 (2016).

58. K. A. Datsenko, B. L. Wanner, One-step inactivation of chromosomal genes in Escherichia coli K-12 using PCR products. Proc. Natl. Acad. Sci. U. S. A. 97, 6640–6645 (2000).

